# The integrated stress response regulates 18S nonfunctional rRNA decay in mammals

**DOI:** 10.1101/2024.07.30.605914

**Authors:** Aaztli R. Coria, Akruti Shah, Mohammad Shafieinouri, Sarah J. Taylor, Wilfried Guiblet, Jennifer T. Miller, Indra Mani Sharma, Colin Chih-Chien Wu

## Abstract

18S nonfunctional rRNA decay (NRD) detects and eliminates translationally nonfunctional 18S rRNA. While this process is critical for ribosome quality control, the mechanisms underlying nonfunctional 18S rRNA turnover remain elusive. NRD was originally identified and has exclusively been studied in *Saccharomyces cerevisiae.* Here, we show that 18S NRD is conserved in mammals. Using genome-wide CRISPR genetic interaction screens, we find that mammalian NRD acts through the integrated stress response (ISR) via GCN2 and ribosomal protein ubiquitination by RNF10. Selective ribosome profiling reveals nonfunctional 18S rRNA induces translational arrest at start sites. Indeed, biochemical analyses demonstrate that ISR activation limits translation initiation and attenuates collisions between scanning 43S preinitiation complexes and nonfunctional 80S ribosomes arrested at start sites. Overall, the ISR promotes nonfunctional 18S rRNA and 40S ribosomal protein turnover by RNF10-mediated ubiquitination. These findings establish a dynamic feedback mechanism by which the GCN2-RNF10 axis surveils ribosome functionality at translation initiation.

## INTRODUCTION

The ribosome is central to controlling gene expression and translating genomic information into functional proteins. This process is governed by the decoding center of the small ribosomal subunit and the peptidyl transferase center of the large ribosomal subunit^1^. In *Saccharomyces cerevisiae*, rRNAs carrying mutations that render the ribosome nonfunctional by impacting the decoding center of 18S rRNA or the peptidyl transferase center of 25S rRNA are downregulated through a late-acting rRNA quality control pathway termed nonfunctional rRNA decay (NRD)^2,3^. NRD targets defective ribosomal subunits for degradation through specific pathways that depend on the site affected by the mutation: 18S NRD eliminates nonfunctional 18S rRNAs and 25S NRD degrades nonfunctional 25S rRNAs^2,3^. A crucial distinction between the two quality control pathways is that ongoing translation is required for 18S NRD but not for 25S NRD^3^. While NRD is well-studied in *S. cerevisiae,* the molecular mechanisms associated with the recognition and degradation of nonfunctional rRNA remain obscure. Additionally, since these quality control pathways have been exclusively studied in budding yeast^2–9^, they remain unexplored in mammals.

In response to diverse cellular stress conditions, cells have evolved quality control mechanisms to detect homeostatic perturbations and elicit appropriate cellular responses. One common response involves the repression of protein synthesis through the phosphorylation of eukaryotic initiation factor 2⍺ (eIF2⍺) by kinases that respond specifically to distinct stresses, known as eIF2⍺ kinases^10^. In mammals, there are four eIF2⍺ kinases— GCN2, PKR, PERK, and HRI. From a canonical perspective, GCN2 is activated by uncharged tRNA during amino acid starvation, PKR by double-stranded RNA during viral infection, PERK by unfolded proteins in the endoplasmic reticulum, and HRI by heme limitation. Activation of each kinase initiates the integrated stress response (ISR), leading to reduction of global translation initiation and upregulation of selected genes, including the essential bZIP transcription factor ATF4, required for downstream stress response pathways to promote cellular recovery^11^. GCN2 is activated by uncharged tRNAs^12–14^. However, recent studies show that GCN2 can also be activated by stalled ribosomes independently of uncharged tRNAs^15,16^ and by ribosome collisions, with disomes as a minimal unit^17–19^. In addition, GCN2 is preferentially activated by collided ribosomes with a vacant A site in the leading ribosome in yeast^18,20^. Biochemical studies demonstrate that GCN2 can be activated by the purified ribosomal P-stalk, composed of uL10 and two heterodimeric P1/P2 proteins^21–23^. In line with these *in vitro* observations, a recent study shows differential requirements for the P-stalk proteins in activating GCN2^24^. While P1/P2 of the P-stalk proteins are dispensable for GCN2 activation by uncharged tRNAs resulting from amino acid starvation, at least one of two distinct P1/P2 heterodimers is required for GCN2 activation induced by amino acid starvation-independent ribosome stalling. These studies highlight the pivotal role of GCN2 in monitoring the status of elongating ribosomes to finetune translation initiation and to maintain cellular homeostasis through the ISR.

When ribosomes stall, a triage process, the ribosome-associated quality control (RQC) pathway, is activated^25–27^. Upon prolonged stalling on mRNAs, trailing ribosomes collide with stalled ribosomes, resulting in the formation of a new interface between the two collided small ribosomal subunits^28^. The E3 ubiquitin ligase, ZNF598 (Hel2 in yeast), recognizes this interface and catalyzes site-specific ubiquitination of ribosomal proteins eS10 (RPS10) and uS10 (RPS20)^29–33^. This ubiquitination is critical for recruiting the factors responsible for cleaving the mRNA^34^ and for splitting the stalled ribosomes into subunits for recycling^30,35^. Resolving ribosome collisions is essential to avoid general stress responses, including the integrated stress response (ISR) and the ribotoxic stress response (RSR)^17^, as well as innate immune signaling^36,37^. Moreover, ubiquitination of ribosomal proteins uS3 (RPS3) and uS5 (RPS2) has been associated with various cellular stressors that abolish translation^38^ and the degradation of 40S ribosomal proteins^39,40^. Specifically, the ubiquitin E3 ligase, RNF10, initiates hierarchical ubiquitination of ribosomal proteins uS3 and uS5 to target stalled single elongating ribosomes and stalled 43S preinitiation complexes (PIC) for elimination^39,40^. Mag2, the yeast homologue of RNF10, has been shown to play a critical role in recognizing nonfunctional 80S ribosomes stalled at initiation codons in recent studies^5,6^, suggesting the existence of 18S NRD in mammals and the possible role of RNF10 in the 18S NRD pathway.

In this study, we developed an orthogonal human 18S rRNA expression system to show the conservation of 18S NRD in mammalian cells. Using fitness-based CRISPR genetic interaction screens, we identified a negative feedback mechanism that regulates translation initiation to facilitate the ubiquitination of ribosomal proteins and hence the turnover of nonfunctional 18S rRNA. Specifically, nonfunctional 18S rRNA leads to prolonged ribosome arrest at start codons across the transcriptome. These decoding-incompetent ribosomes provoke the ISR through GCN2 to reduce translation initiation, thereby preventing collisions between scanning 43S PICs and decoding-incompetent ribosomes. This negative feedback loop ensures that RNF10 can target decoding-incompetent ribosomes for destruction. These findings highlight the role of the ISR and its kinase, GCN2, as critical upstream regulators in the turnover of the 40S ribosomal subunit.

## RESULTS

### Conservation of 18S nonfunctional rRNA decay in human cells

To systematically explore how mammalian cells cope with nonfunctional 18S rRNA, we developed an orthogonal human 18S rRNA expression system in which 18S rDNA is flanked by a human 5’ external transcribed spacer (5’ ETS) and internal transcribed spacer 1 (ITS1) to ensure proper 18S rRNA maturation^41^ (Figure 1A). We used an RNA polymerase I promoter and terminator to achieve high levels of transcription. At the tip of helix 9 near the 5ʹ end of the orthogonal 18S rRNA, we inserted either a tandem MS2 hairpin (dubbed 18S-tMS2) for ribosome isolation^5^ or a hybridization hairpin (dubbed 18S-H) for detection by Northern blotting^41^ (Figure S1A). Helix 9 has been previously reported as a segment that can harbor an exogenous tag without disrupting its tertiary structure within the rRNA^42^. To confirm that helix 9 is properly folded despite these insertions, we used *in vivo* SHAPE (<u>s</u>elective 2ʹ-<u>h</u>ydroxyl <u>a</u>cylation analyzed by <u>p</u>rimer <u>e</u>xtension) RNA structure probing to examine the local secondary structure of helix 9 and the inserted hairpin. We treated HeLa cells expressing either the tMS2- or H-containing orthogonal 18S rRNA with the SHAPE reagent, which probes nucleotide flexibility^43,44^. We then sequenced and analyzed the amplicons derived from tMS2- or H-tagged helix 9 using a high-throughput mutational profiling (MaP) procedure^45^. SHAPE-derived secondary structures validated that both hairpin insertions extend from the tip of the 18S helix 9 stem-loop without disrupting its folding *in vivo* (Figures 1B and S1B-D).

**Figure 1.**
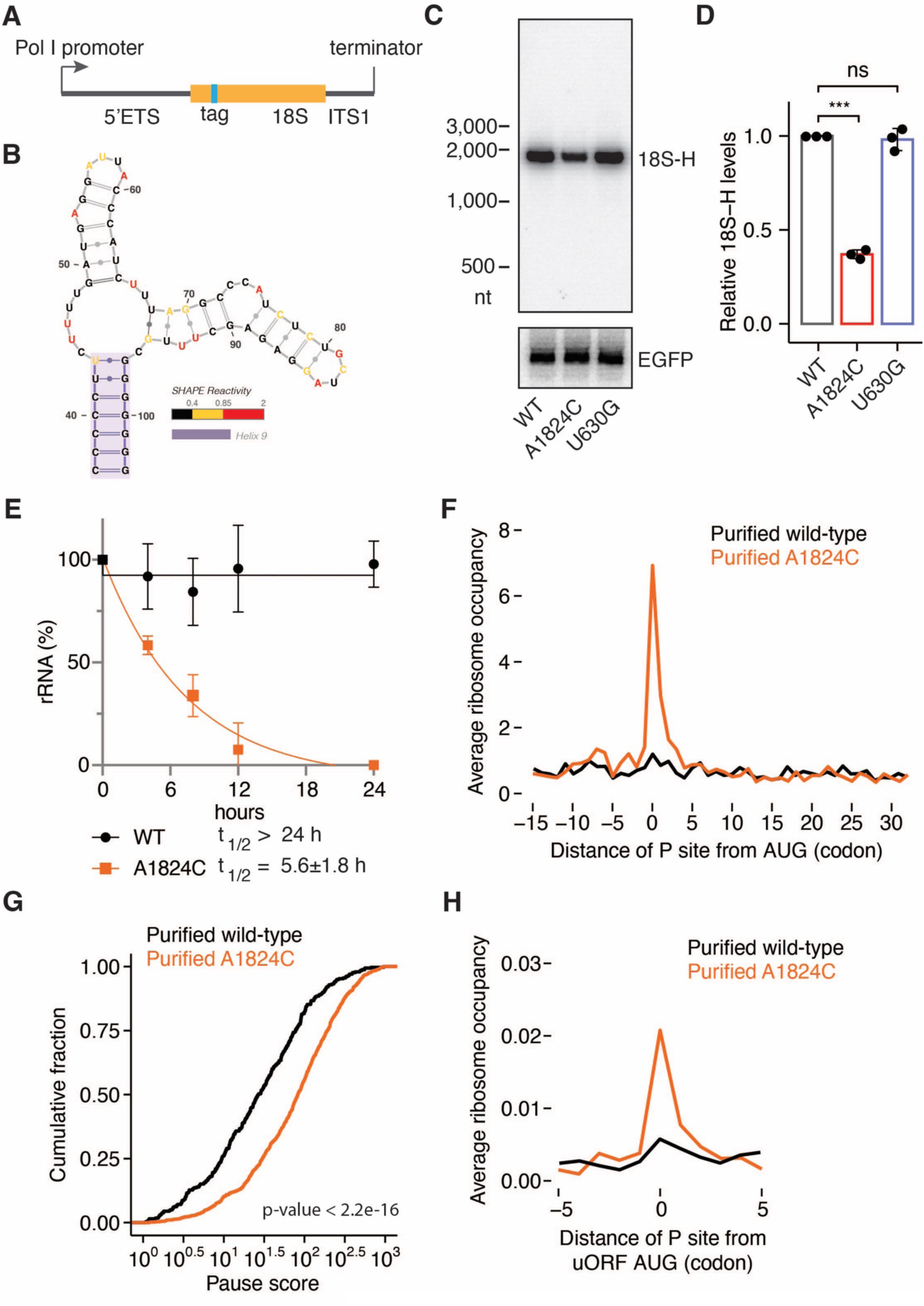
An orthogonal human 18S rRNA expression system. **(A)** Schematic depicting strategy used to express human 18S rRNA. The orthogonal 18S expression construct contains a Pol I promoter, 5’ external transcribed spacer (5ʹ ETS), 18S rRNA sequence including a tag (indicated in cyan), and internal transcribed spacer 1 (ITS1). The tag was introduced into 18S rRNA at helix 9 (also see Figure S1A). **(B)** A SHAPE-derived minimum free energy secondary structure of helix 9, including a hybridization (H) tag. Nucleotides exhibiting high (2- 0.85), medium (0.85- 0.4), and low (0- 0.4) SHAPE reactivity are color-coded in red, yellow, and black, respectively. **(C)** Northern blot analysis for steady-state levels of 18S rRNA carrying indicated mutation, using a probe against H-tag. WT: wild-type. A1824C: decoding center mutation. U630G: a benign mutation. EGFP was used as a transfection and loading control. Also see Figure S1F. **(D)** Quantification of Northern blots from (C) measuring steady-state levels of 18S:WT, A1824C and U630G relative to EGFP levels (n= 3). WT replicates were set to 1. Error bars denote the standard deviation. Significance was calculated using unpaired Student’s t-test (***: p < 0.001, ns: not significant). **(E)** Half-life measurement of 18S:WT (black) and A1824C rRNA (orange). RNA was harvested from cells that constitutively express 18S:WT or A1824C rRNA (see Figure S1H) at 0, 6, 12, 18, and 24 hours after Pol I transcriptional inhibition. 18S rRNA levels were quantified from triplicates and levels at 0 hour were set to 100%. Error bars represent the standard deviation (SD). Half-lives are denoted with ± SD. **(F)** Average ribosome occupancy aligned at start codons from affinity-purified (see Figure S2B) WT (black trace) or decoding-incompetent ribosomes (A1824C). Ribosomal P sites of footprints are plotted relative to the translation start site. The plot represents one biological replicate (n= 3). **(G)** Cumulative frequency histogram of pause scores at start codons shows enhanced stalling of purified decoding-incompetent ribosomes (A1824C, orange trace) compared to wild-type ribosomes (black trace). **(H)** Metagene analysis of average ribosome occupancy aligned at annotated uORF start codons from affinity-purified wild-type (black trace) and decoding-incompetent ribosomes (A1824C, orange trace). Offsets were set to show the ribosomal P site of the footprints.

mRNA decoding by the ribosome involves the coordination of the universally conserved monitoring bases— G626, A1824, and A1825 (human numbering; G530, A1492 and A1493 in *E. coli*) in the decoding center of the 40S ribosomal subunit (Figure S1E) to facilitate codon-anticodon interactions^1,46–48^. Mutations at the equivalent of human A1824 in the 18S rRNA trigger 18S NRD in *S. cerevisiae*^2,3^. While NRD is not observed in bacteria^49,50^, whether such turnover of nonfunctional 18S rRNA is conserved in mammals has not yet been tested. To address this, we introduced a single nucleotide mutation in the decoding center of the orthogonal 18S rRNA— specifically, A1824C (Figure S1E). After transfecting the expression constructs into HeLa cells, Northern blotting and primer extension experiments revealed a ∼2.5-fold reduction in 18S:A1824C rRNA levels compared to 18S:WT rRNA (Figures 1C, 1D and S1F). In addition, the benign mutation (U630G) that is located close to the decoding center but plays no role in decoding^2^ showed steady-state levels similar to 18S:WT. Indeed, 18S:A1824C rRNA was consistently downregulated across multiple human cell lines, including HEK293 and HaCaT (a human keratinocyte cell line) (Figure S1G).

To determine whether the observed difference at steady state between WT and A1824C 18S rRNA resulted from decay rather than synthesis, we engineered HeLa cells that constitutively express the orthogonal 18S rRNA constructs and determined the levels of 18S:WT and 18S:A1824C rRNA to be approximately ∼8% and ∼3% of the endogenous 18S rRNA, respectively (Figure S1H). We then performed decay kinetics by halting transcription with the RNA polymerase I inhibitor (CX-5461) and monitored 18S rRNA levels over a 24-hour period. 18S:A1824C rRNA indeed underwent faster turnover (t_1/2_= 5.6 ± 1.8 h) than 18S:WT rRNA which remained stable over the experimental timeframe (t_1/2_> 24 h) (Figure 1E). The long half-life of the control is consistent with the known stability of mature wild type 18S rRNA^51^. Taken together, these findings provide compelling evidence for the conservation of 18S NRD in mammalian cells.

### Decoding-incompetent ribosomes are predominantly arrested at translation start sites

Having established that 18S NRD is well-conserved in human cells, we next asked whether nonfunctional 18S rRNA can be assembled into functional 40S subunits and actively translating ribosomes. Polysome profiling followed by primer extension analysis revealed deep sedimentation of 18S:WT rRNA in the 40S, 80S, and polysome fractions (fractions 6-10). In contrast, 18S:A1824C rRNA sedimented in the 40S, 80S, and light polysome fractions (fractions 5-7) (Figure S2A). These sedimentation profiles are consistent with the notion that the mutant 18S rRNA is incorporated into fully mature 40S subunits and 80S ribosomes similar to budding yeast^2^.

To gain insights into how decoding-incompetent ribosomes impact global translation, we performed selective ribosome profiling (Figure S2B). After RNase digestion and isolation of tMS2-tagged ribosomes by affinity purification (Figure S2C), we prepared footprint libraries from purified wild-type and decoding-incompetent ribosomes. Because the commonly used ribonuclease, RNase I, digests the inserted tMS2 tag, we chose nuclease P1 for sample preparation^52^. As a result, the average length of ribosome footprints is ∼35 nt (Figure S2D), larger than those seen for samples prepared with RNase I. We observed strong enrichment of footprints from purified mutant ribosomes at start codons throughout the transcriptome including, for instance, HIST1H1C and ZMAT2 mRNAs (Figure S2E). We calculated the average ribosome occupancy at the start codons of all transcripts and found that mutant ribosome footprints were ∼7-fold enriched in this region compared to wild-type ribosome footprints (Figure 1F). Moreover, when computing a pause score for each start codon, the median pause score increased by ∼3-fold in the mutant ribosomes compared to wild-type ribosomes (Figure 1G), emphasizing the transcriptome-wide nature of this effect. This perturbation skewed the distribution of the mutant ribosome footprints toward the 5ʹ end of the open reading frames (ORFs) (Figure S2F), as assessed by computing a polarity score for each gene (see Methods)^53^. In addition to the start codons of main ORFs, mutant ribosome footprints are also enriched (∼4-fold) at the start codons of upstream ORFs (uORFs) (Figure 1H). Together, these results indicate that 18S:A1824C rRNA is incorporated into mature 80S ribosomes, resulting in decoding-incompetent ribosomes that stall, predominantly at start codons, across the transcriptome.

### A fitness-based CRISPR screen identifies GCN2 as an NRD component

Upon integrating the orthogonal 18S rRNA into cells, we noted that cells expressing 18S:A1824C rRNA (dubbed 18S:A1824C cells) consistently showed reduced cell fitness compared to those expressing 18S:WT rRNA (dubbed 18S:WT cells) (Figure 2A). This decrease occurs despite the relatively low abundance of 18S:A1824C (Figure S1H). We hypothesized that the difference in growth rate is likely due to the presence of nonfunctional 18S rRNA. Therefore, we took advantage of this cell fitness phenotype and performed a genome-wide CRISPR/Cas9 screen^54^ to systematically identify genes associated with the cellular response to nonfunctional 18S rRNA. We transduced 18S:WT and 18S:A1824C cells with a library of single-guide RNAs (sgRNAs) targeting each known protein-coding gene (19,114 genes). After culturing each population for 10 doublings, we deep-sequenced the remaining sgRNAs at four time points (0, 5, 7, and 10 doublings) (Figure 2B).

**Figure 2.**
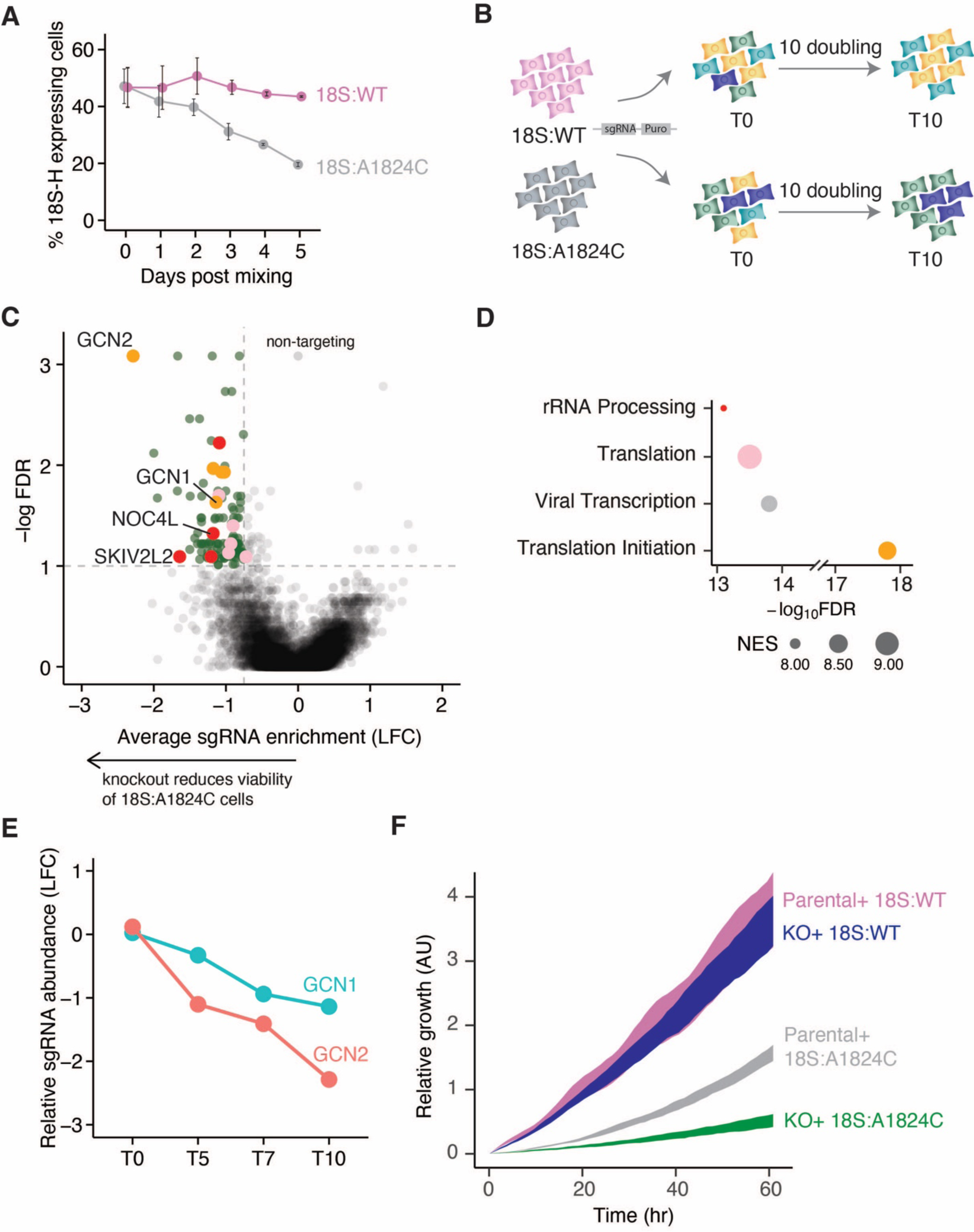
CRISPR screen to identify genes linked to nonfunctional 18S rRNA turnover. **(A)** Competition assays using HeLa cells expressing 18S:WT (magenta) or 18S:A1824C rRNA (grey). 18S:WT or 18S:A1824C cells were co-cultured with non-fluorescent parental HeLa cells and the ratios were determined by flow cytometry at the indicated time points (n= 3). **(B)** Schematic depicting the fitness-based CRISPR knockout screen. The library was introduced into cells expressing either 18S:WT or 18S:A1824C rRNA. Cells were passaged over 10 doublings. **(C)** Volcano plot showing average sgRNA enrichment (log_2_ fold change [LFC], x axis) versus false-discovery rate (FDR) (-logFDR, y axis) at T10. Significant genes (LFC< 0.75 and FDR< 0.1) are highlighted in green. Dot color corresponds to the enriched Gene Ontology (GO) terms in (D). Also see Figures S3A and S3B. **(D)** Top 4 enriched biological processes from T10 in (C). Normalized enrichment score (NES) is denoted by dot size. **(E)** Quantification of the relative abundance (LFC in 18S:A1824C/LFC in 18S:WT) of GCN1 (green) and GCN2 (salmon) sgRNAs. **(F)** Quantification of cell growth by xCELLigence over 66 hours (x axis). Growth phenotype was compared across four cell lines: parental HeLa cells expressing 18S:WT (magenta) or 18S:A1824C rRNA (grey) and GCN2KO cells expressing 18S:WT (blue) or 18S:A1824C rRNA (green). Curve width is representative of the standard deviation over 4 replicates.

While overall trends identified at these time points were similar, depletion of essential genes was more pronounced at the later time points (Figure S3A), indicative of the effectiveness of CRISPR/Cas9-mediated gene editing. When comparing the changes in sgRNA abundance between 18S:WT and 18S:A1824C cells after 10 doublings, we focused on genes whose loss-of-function led to further impediment of cell fitness. A total of 116 genes were significantly depleted in the 18S:A1824C population (Figure 2C), and gene ontology (GO) analysis revealed strong enrichment for genes implicated in translation, rRNA processing, and translation initiation (Figure 2D). Genes involved in 18S rRNA processing and maturation, including SKIV2L2, RRP7, RRP9, and NOC4L^55^, were among those identified (Figures 2C and S3B). Moreover, the screen also identified HBS1L (Figure S3B), whose yeast homologue is an established 18S NRD factor^4^. Indeed, short hairpin RNA (shRNA)-mediated knockdown of HBS1L resulted in accumulation of 18S:A1824C rRNA (Figure S3C).

In addition to these factors, the screen revealed several genes with no known 18S NRD-related function. One of the four mammalian eIF2⍺ kinases— GCN2 emerged as the most prominent hit and its co-activator, GCN1, was also significantly de-enriched in 18S:A1824C cells (Figures 2C, 2E and S3D). Because GCN2 has not been implicated previously in cellular responses to nonfunctional 18S rRNA, we chose to focus on this gene. We validated the cell growth phenotypes by knocking out GCN2 in 18S:WT and 18S:A1824C cells. Consistent with the CRISPR screen results, loss of GCN2 exacerbated the moderate growth defects in 18S:A1824C cells though it had no discernible effect on 18S:WT expressing cells (Figure 2F).

### GCN2 and its effectors are required for 18S NRD

Given the hypothesis that nonfunctional 18S rRNA reduces cell fitness and this effect is exacerbated by loss of GCN2 (Figure 2F), we predicted that GCN2 plays a role in 18S NRD and its ablation might stabilize 18S:A1824C rRNA. Strikingly, we observed a marked ∼3-fold increase in 18S:A1824C rRNA levels in cells depleted of GCN2 by shRNA (Figures 3A and 3B) or CRISPR/Cas9 (Figure S4A) while 18S:WT rRNA levels were unaffected. We further found that GCN2 knockout (KO) cells complemented with wild-type GCN2, but not a catalytically inactive mutant (D848N), are competent to degrade 18S:A1824C rRNA (Figures 3C and 3D), implying that GCN2 kinase activity is critical for 18S NRD. In line with this observation, knockdown of GCN1 also stabilized 18S:A1824C rRNA levels (Figures S4B and S4C), in agreement with our CRISPR screen results (Figures 2C and 2E). These results are readily reconciled with the observation that the fitness of 18S:A1824C cells was further impeded upon GCN2 knockout (Figure 2F).

**Figure 3.**
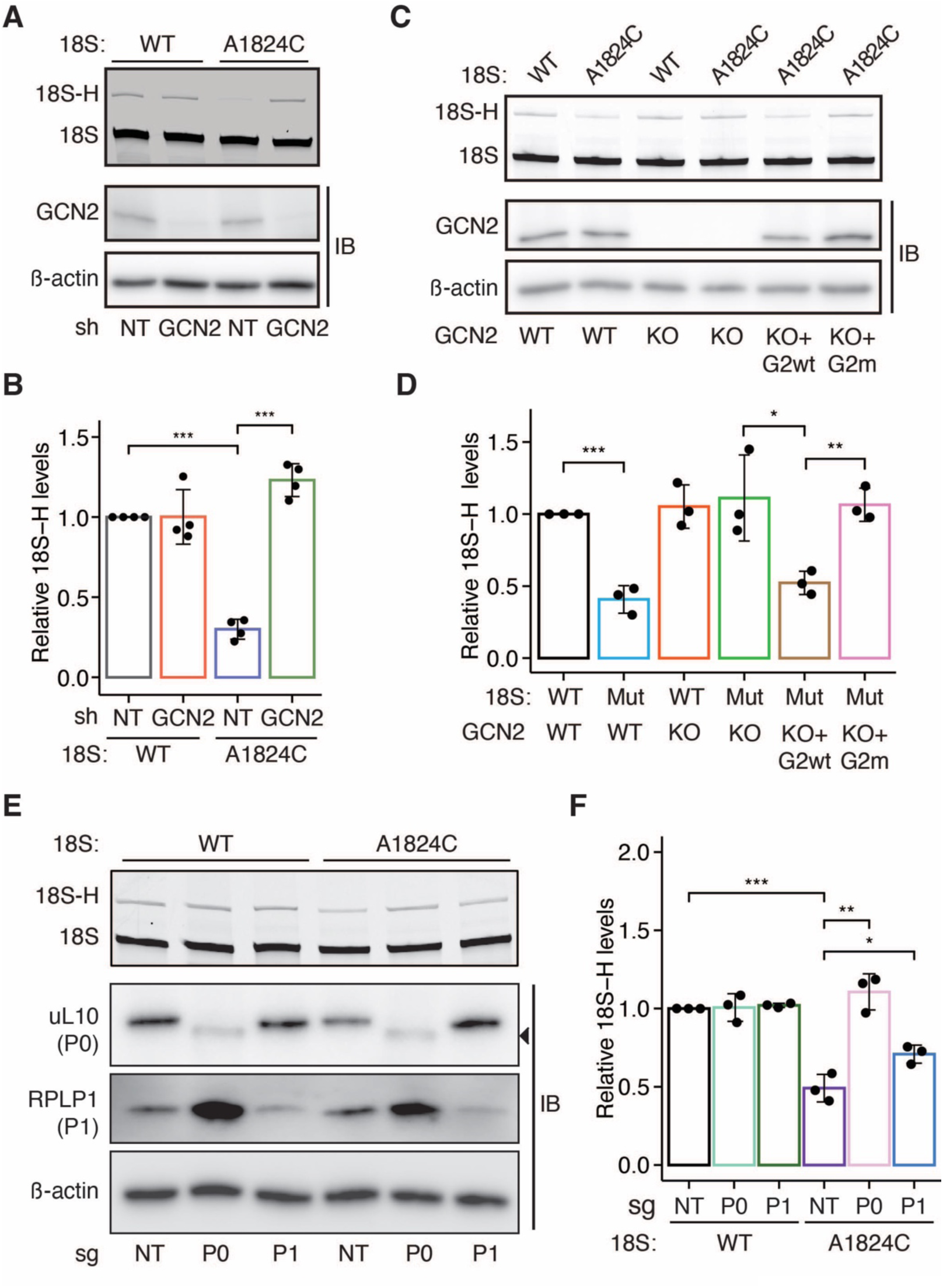
GCN2 and its activators are required for nonfunctional 18S rRNA turnover. **(A)** Primer extension analysis of orthogonal 18S rRNA (WT or A1824C) upon shRNA-mediated knockdown of GCN2. The top band is derived from the orthogonal 18S-H rRNA, and the bottom band is endogenous 18S rRNA. Bottom: immunoblots for GCN2 protein levels. β-actin serves as a loading control. NT: non-targeting. **(B)** Quantification of primer extension assay shown in (A) (n= 4). **(C)** Primer extension analysis of 18S rRNA in parental HeLa or GCN2KO cells complemented with WT GCN2 (G2wt) or catalytically inactive GCN2 (D848N, G2m). Bottom: immunoblots for GCN2 expression levels. **(D)** Quantification of primer extension assays shown in (C) (n= 3). Mut: A1824C. **(E)** Primer extension analysis of 18S:WT or A1824C upon CRISPR/Cas9-mediated editing of uL10 (P0) or RPLP1 (P1). Bottom: Control immunoblots for uL10, RPLP1, and β-actin. Arrowhead indicates truncated uL10 resulting from Cas9-mediated gene editing (also see Figure S4D). **(F)** Quantification of 18S-H levels upon depletion of uL10 or RPLP1 in 18S:WT or A1824C cells shown in (E) (n= 3). In (B), (D), and (F), error bars denote the standard deviations and significance was calculated using unpaired Student’s t-test (*: p < 0.05, **: p < 0.01, and ***: p < 0.001).

There are multiple lines of evidence arguing that the ribosomal P-stalk, positioned near the A site, is indispensable for GCN2-mediated eIF2⍺ phosphorylation both *in vitro* and *in vivo*^21,22,24^, particularly during amino acid starvation-independent ribosome stalling^24^. Given that both GCN2 and GCN1 are essential for 18S NRD, we wondered whether the ribosomal P-stalk might also play a role in 18S NRD. Using CRISPR/Cas9, we ablated the ribosomal P-stalk genes RPLP0 and RPLP1, which encode uL10 and P1, respectively. While RPLP0 ablation increased 18S:A1824C rRNA to levels comparable to 18S:WT, RPLP1 ablation showed a less pronounced effect (Figures 3E and 3F). This is likely because sgRNA-mediated editing deletes the C-terminal helical spine of uL10, leading to disintegration of the ribosomal P-stalk protrusion composed of uL10 C-terminus and two heterodimeric P1 and P2 proteins (Figure S4D). These data are consistent with previous structural, genetic, and biochemical evidence showing direct association/binding of the GCN complex with the P-stalk proteins^21,22,56^. Collectively, we find that ablation of GCN2 kinase activity and co-factors required for GCN2 activation consistently reduces 18S NRD activity, indicating that eIF2⍺ phosphorylation by GCN2 is critical for 18S NRD. Of the four eIF2⍺ kinases, GCN2 is closely related to protein synthesis unlike PERK, PKR, and HRI, making this newly discovered role in regulating nonfunctional rRNA turnover reasonable.

### Decoding-incompetent ribosomes trigger GCN2-dependent eIF2⍺ phosphorylation

GCN2 regulates global translation initiation by phosphorylating eIF2⍺ at serine 51 in response to various cellular stressors^57^. In light of recent studies proposing ribosome-centric mechanisms that activate GCN2 independently of uncharged-tRNA^15,17^, we asked whether decoding-incompetent ribosomes trigger eIF2⍺ phosphorylation since they are found among actively translating ribosomes (Figures S2A and 1F-H). Using immunoblotting, we observed a ∼2-fold increase in eIF2⍺ phosphorylation levels and a strong concomitant induction of ATF4 translation in 18S:A1824C-transfected cells, indicating that the ISR is activated (Figure 4A). Importantly, we found that 18S:A1824C rRNA did not induce eIF2⍺ phosphorylation and ATF4 expression in GCN2KO and GCN1KO cells (Figures 4A and 4B), consistent with recent findings that GCN1 is essential for GCN2-dependent eIF2⍺ phosphorylation in an amino acid starvation-independent manner^18,24^.

**Figure 4.**
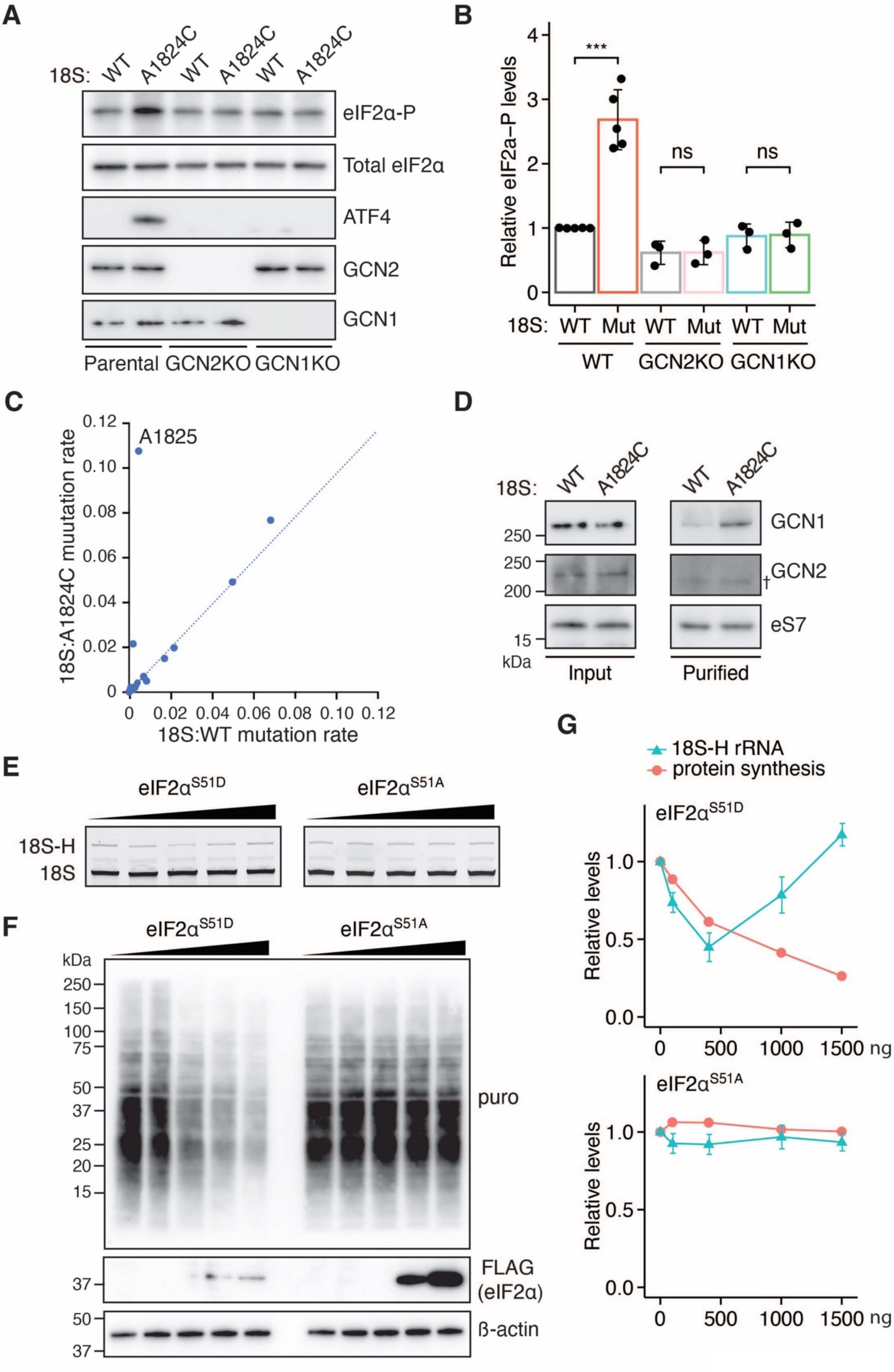
GCN2 regulates 18S NRD by finetuning eIF2⍺ phosphorylation. **(A)** Immunoblots for eIF2⍺ phosphorylation (eIF2⍺-P), total eIF2⍺, and ATF4 levels in parental HeLa, GCN2KO, and GCN1KO cells transfected with 18S:WT or 18S:A1824C rRNA expression construct. Total eIF2⍺ serves as a loading control. **(B)** Quantification of eIF2⍺ phosphorylation levels shown in (A) (n ≥ 3). Relative eIF2⍺-P levels were calculated by taking the ratio of total eIF2⍺ and eIF2⍺-P. Error bars denote the standard deviation. Significance was calculated using unpaired Student’s t-test (***: p < 0.001). **(C)** Scatter plot comparing DMS mutation rates of nucleotides located within the A site of wild-type (x-axis) and decoding-incompetent (A1824C, y-axis) ribosomes. Also see Figure S5A. **(D)** Immunoblotting analysis of affinity-purified wild-type and decoding-incompetent (A1824C) ribosomes with indicated antibodies. Ribosomal protein eS7 serves as a loading control. † denotes a non-specific band recognized by the antibody. **(E)** Primer extension analysis of 18S:A1824C rRNA levels in GCN2KO cells upon titrating a plasmid encoding phosphomimetic eIF2⍺^S51D^ (left) or non-phosphorylatable eIF2⍺^S51A^ (right). **(F)** Immunoblots for protein synthesis (nascent polypeptide chains released by puromycin incorporation) and expression levels of eIF2⍺^S51D^ or eIF2⍺^S51A^ from cell lysates shown in (E). β- actin is a loading control. **(G)** Quantification of 18S:A1824C rRNA levels (salmon) and protein synthesis (green) in cells titrated with eIF2⍺^S51D^ (top) or eIF2⍺^S51A^ (bottom).

We next asked whether decoding-incompetent ribosomes derived from 18S:A1824C rRNA contain a vacant A site in the cell, consequently activating GCN2. To this end, we turned to RNA structure probing to examine difference in the accessibility of rRNA nucleotides in the decoding center. We treated cells expressing tMS2-tagged 18S:WT or A1824C with dimethyl sulfate (DMS), which probes the accessibility of the Watson-Crick base-pairing interface of adenosine and cytosine. Aminoacyl-tRNA binding to the ribosomal A site shields nucleotides A1824 and A1825 (A1492 and A1493 in *E. coli*) in the 18S rRNA from DMS modification^58^. Quantifying the accessibility of these nucleotides can thereby assess whether the A site is vacant. After isolation of tagged ribosomes, we designed an amplicon derived from the decoding center and found a ∼27-fold increase in the mutation rate (i.e. more accessible) at position A1825 in samples from decoding-incompetent ribosomes relative to those from wild-type ribosomes (Figures 4C and S5A). These structural probing data indicate that an inactivating mutation in the decoding center prevents aminoacyl-tRNA binding in the A site.

Considering the binding capacity of GCN1/20 for the ribosome^59^, we hypothesized that GCN1/20 may be recruited to decoding-incompetent ribosomes. Purification of wild-type and decoding-incompetent ribosomes followed by immunoblotting showed that GCN1 preferentially binds to mutant ribosomes compared to wild-type ribosomes (Figure 4D). While we did not observe direct binding of GCN2 to decoding-incompetent ribosomes, this association may be transient and/or compromised by steps such as RNase treatment during sample preparation. Taken together, these data argue that arrested decoding-incompetent ribosomes with a vacant A site activate the GCN2-dependent ISR.

### GCN2 ensures 18S NRD by finetuning translation initiation

Our data point to a model where decoding-incompetent ribosomes trigger GCN2-dependent eIF2⍺ phosphorylation to reduce ribosome loading onto mRNAs, thereby attenuating translation initiation to facilitate 18S NRD. We therefore evaluated whether eIF2⍺ phosphorylation could complement loss of GCN2 and promote 18S NRD by titrating a plasmid encoding phosphomimetic eIF2⍺^S51D^ into GCN2KO cells expressing 18S:A1824C rRNA. In contrast to chemical and environmental stressors that can trigger eIF2⍺ phosphorylation, the ectopic expression of phosphomimetic eIF2⍺^S51D^ allows us to examine the direct effect of eIF2⍺ phosphorylation on nonfunctional 18S rRNA turnover without inducing unwanted cellular stress responses. Strikingly, we found that intermediate, rather than high, levels of phosphomimetic eIF2⍺^S51D^ complemented the loss of GCN2 and promoted 18S NRD whereas titration of a plasmid encoding non-phosphorylatable eIF2⍺^S51A^ had no effect (Figures 4E and 4G). Protein synthesis, as measured by puromycin incorporation, decreased drastically in eIF2⍺^S51D^-expressing cells at the high end of the titration curve, reflecting the anticipated reduction in global translation initiation associated with eIF2⍺ phosphorylation (Figure 4F). In contrast, titration of non-phosphorylatable eIF2⍺^S51A^ resulted in no significant changes in protein synthesis even though the expression levels of eIF2⍺^S51A^ were higher than those of eIF2⍺^S51D^. These results reveal a “V-shaped” dose-response curve of 18S NRD activity in response to eIF2⍺ ^S51D^ levels (Figure 4G), suggesting that an optimal regime of translation initiation is critical for the turnover of nonfunctional 18S rRNA. Additionally, this dependence on eIF2⍺ phosphorylation is consistent with the kinase activity of GCN2 being required for degrading nonfunctional 18S rRNA (Figures 3C and 3D). Taken together, our results suggest that upon sensing the vacant A site of arrested decoding-incompetent ribosomes the GCN complex maintains an optimal level of translation initiation for 18S NRD through finetuning eIF2⍺ phosphorylation.

### Loss of GCN2 exacerbates start codon arrest resulting in 43S-80S collisions

eIF2⍺ phosphorylation by eIF2⍺ kinases reduces translation initiation and subsequent ribosome loading onto mRNAs. If GCN2 promotes the degradation of 18S A1824C rRNA through eIF2⍺ phosphorylation, we reasoned that GCN2 ablation would lead to an accumulation of decoding-incompetent ribosomes at start codons. To test this prediction, we performed selective ribosome profiling with tMS2-tagged wild-type and decoding-incompetent ribosomes in parental HeLa cells and GCN2KO cells. Metagene analysis showed ∼2-fold enrichment of mutant ribosomes at the start codons of main ORFs (Figures 5A-5B) and uORFs (Figure S5B) in GCN2KO cells compared to parental HeLa cells while there is no discernible enrichment of wild-type ribosomes. Additionally, we readily observed a moderate increase in the start codon peak in GCN2-ablated cells expressing 18S:A1824C rRNA even without enriching tagged ribosomes by affinity purification (Figure S5B), indicative of a mild perturbation in the translational landscape by decoding-incompetent ribosomes despite their low abundance.

**Figure 5.**
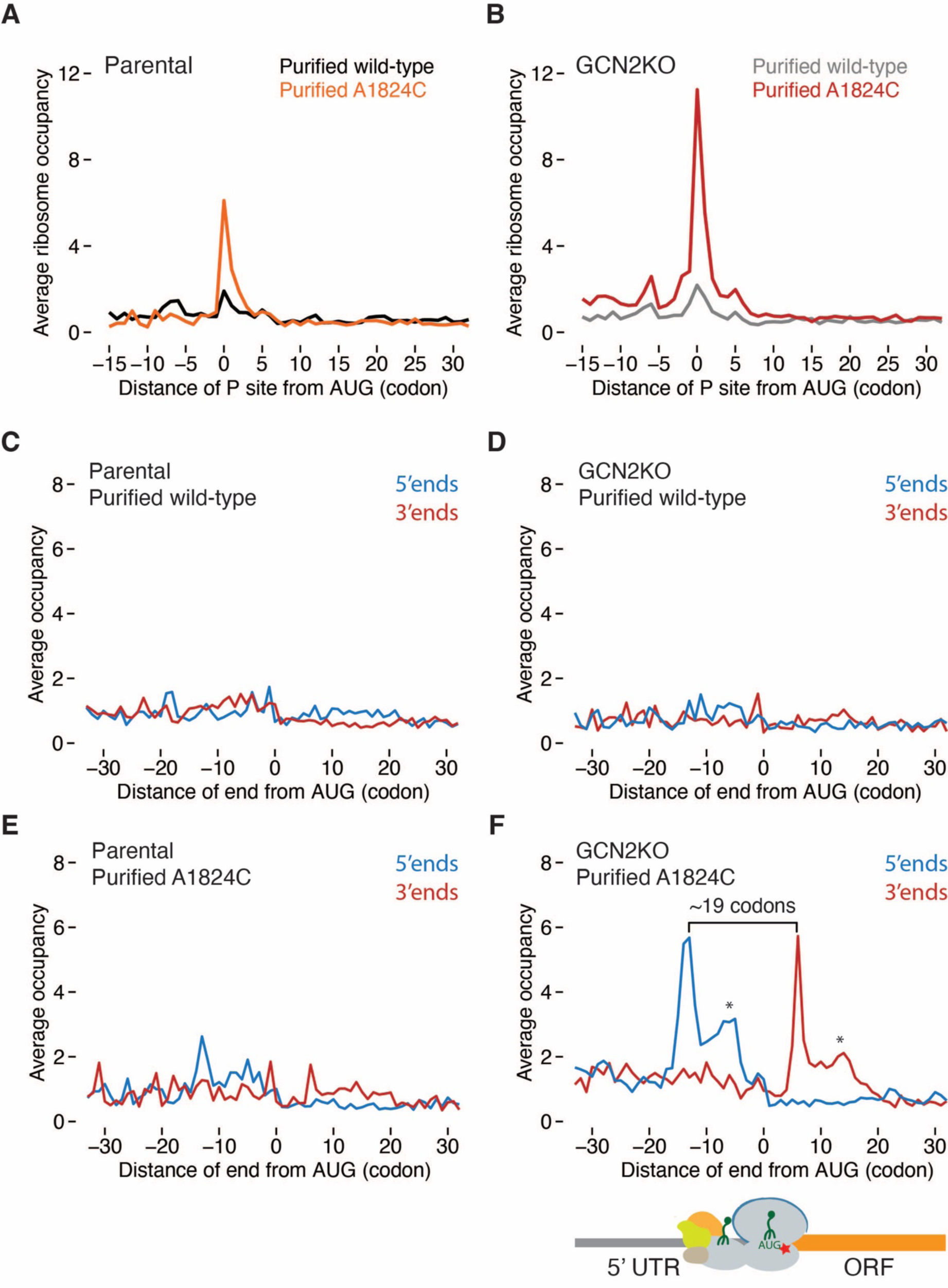
Loss of GCN2 exacerbates start codon arrest resulting in 43S-80S collisions. **(A)** Average ribosome occupancy aligned at start codons from affinity-purified wild-type ribosomes (black trace) and decoding-incompetent ribosomes (A1824C, orange trace) centered at the ribosomal P site. **(B)** Average ribosome occupancy aligned at start codons in GCN2KO cells from affinity-purified wild-type ribosomes (grey trace) and A1824C ribosomes (red trace) centered at the ribosomal P site. **(C – F)** Metagene analysis at start codons using the 5ʹ (blue) or 3ʹ ends (red) of affinity-purified 43S-80S footprints from wild-type ribosomes in parental HeLa (C) and GCN2KO cells (D) or from decoding-incompetent ribosomes in parental HeLa (E) and GCN2KO cells (F) with schematic depicting collisions between a 43S PIC and an 80S ribosome. There is strong enrichment of 43S-80S footprints centered around start codons in samples derived from 18S:A1824C in GCN2KO cells (F). The distance between the 5ʹ end and 3ʹ end of the main peak is denoted. Loss of GCN2 enhances 43S-80S footprints around the start codon. Asterisks indicate 43S-80S footprints downstream the start codon. Red star denotes A1824C mutation in the decoding center of the small subunit.

Although we previously showed that collided ribosomes during translation elongation could activate the ISR through GCN2 and the RSR through mitogen-activated triple kinase (MAPKKK), ZAK⍺^17^, it seems unlikely that decoding-incompetent ribosomes arrested at start codons would result in ribosome collisions (i.e. 80S-80S collisions) in the 5ʹ UTRs because subunit joining only occurs when the scanning 43S preinitiation complex (PIC) reaches a start codon^60^. In line with this prediction, activation of ZAK⍺ (monitored through its phosphorylation on Phos-tag gels) was not detected in GCN2KO cells expressing 18S:A1824C rRNA compared to cells treated with an intermediate dose of anisomycin that provokes ribosome collisions (Figure S5D). In addition, p38 phosphorylation, the downstream amplified signal output of ZAK⍺ cascade, was absent in the 18S:A1824C-expressing cells in contrast to anisomycin-stimulated cells (Figure S5D). We further verified the absence of 80S-80S ribosome collisions by carrying out disome profiling in GCN2-ablated cells that express 18S:WT or A1824C rRNA. In contrast to the ribosome profiling results where we observed an appreciable level of monosome footprints at start codons (Figure S5C), there was no apparent accumulation of disome footprints at the start codons in the metagene analysis (Figure S5E). Altogether, these results indicate that expression of 18S:A1824C rRNA does not induce 80S-80S ribosome collisions.

Given that 80S-80S ribosome collisions are unlikely to take place at start codons, we hypothesized that a scanning 43S PIC may collide with an arrested decoding-incompetent 80S ribosome, thereby dampening nonfunctional 18S rRNA turnover. To directly monitor whether such collisions occur during initiation (i.e. 43S-80S collisions), we purified wild-type and decoding-incompetent ribosomes from parental HeLa and GCN2KO cells, and size-selected fragments 40-80 nt in length to include 43S-80S footprints^52^. Metagene analysis using the 5ʹ or 3ʹ ends of the isolated wild-type ribosome footprints reveals no noticeable enrichment around start codons in either parental (Figure 5C) or GCN2KO cells (Figure 5D). In contrast, footprints from purified decoding-incompetent ribosomes showed strong enrichment at the start codon in GCN2KO cells compared to parental cells (Figures 5E and 5F). The distance between the 5ʹ and 3ʹ ends of the isolated footprints is ∼19 codons (57 nt) (Figure 5F), corresponding to the footprint length protected by 43S-80S collision complexes as reported recently in budding yeast^52^. These observations suggest that 43S PICs queue behind decoding-incompetent ribosomes at translation start sites. Curiously, we noted that there are 43S-80S footprints downstream of the start codon in GCN2KO cells (Figure 5F, asterisks), suggesting that a fraction of the 43S-80S collision complexes might be bound by proteins, thereby generating variable footprint lengths. Collectively, these results support the hypothesis that scanning 43S PICs collide with decoding-incompetent ribosomes and that GCN2 curbs translation initiation to alleviate such collisions during the initiation phase of translation.

In principle, leaky scanning at the stop codons of uORFs could allow a 40S ribosomal subunit to continue scanning and collide with a decoding-incompetent ribosome arrested at the start codons of main ORFs, resulting in initiation collisions. To assess whether such a scenario occurs, we computationally excluded transcripts with annotated uORFs^61^ from the metagene analysis. The presence or absence of uORF-containing transcripts had no impact on the pause amplitude at the start codon (Figures S6A-6B), suggesting that the observed initiation collisions are pervasive in GCN2KO cells. Our data highlight the critical role of GCN2 in controlling ribosome loading to reduce collisions during not only defective elongation (80S-80S) but also defective initiation (43S-80S).

### A Genetic Interaction Screen Identifies RNF10 as an NRD Component That Is Epistatic to GCN2

Having established the dependence of 18S NRD on GCN-mediated eIF2⍺ phosphorylation, we next sought to identify the molecular signatures downstream of GCN2 that trigger nonfunctional 18S rRNA turnover. As described above, we carried out a fitness-based CRISPR genetic interaction screen using GCN2KO cells expressing 18S:WT or A1824C rRNA (Figure S4A). Because GCN2 ablation further exacerbated the growth defects imparted by 18S:A1824C rRNA (Figure 2F), we aimed to search for components that could buffer the growth defects in combination with loss of GCN2 (i.e., greater fitness than expected^62^) (Figure S6C). Therefore, we focused on genes whose sgRNAs were enriched after 10 doubling in 18S:A1824C rRNA-expressing GCN2KO cells, indicating that they are dispensable. These genes, when lost, would not result in further decay of 18S:A1824C rRNA. Given that genetic interactions are rare, we identified 17 genes, with RNF10, an E3 ubiquitin ligase, being one of the top genetic interactors of GCN2 in response to nonfunctional 18S rRNA (Figure 6A).

**Figure 6.**
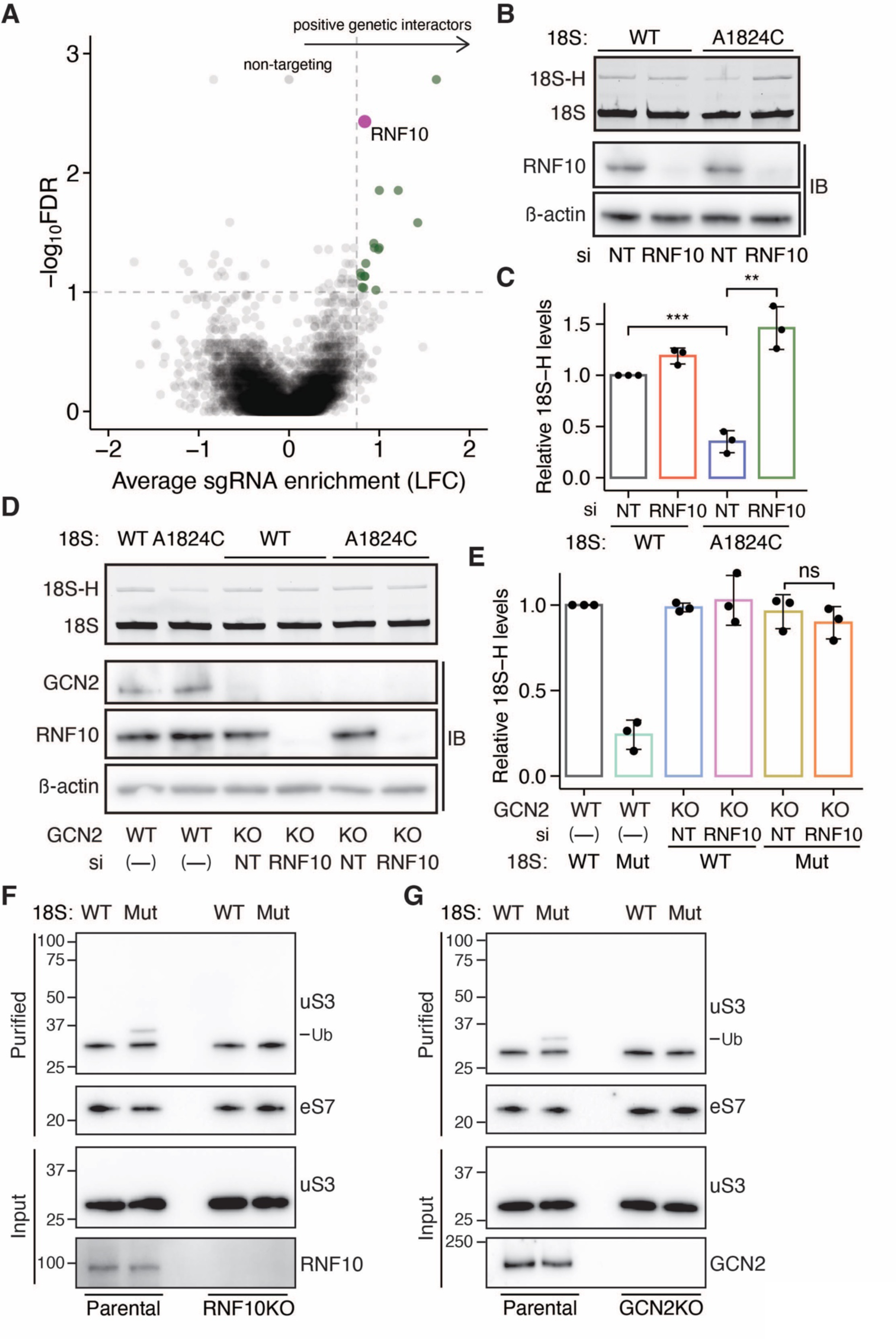
Identification of a digenic interaction between GCN2 and RNF10. **(A)** Volcano plot showing average sgRNA enrichment (log_2_ fold change [LFC], x axis) versus false-discovery rate (FDR) (-logFDR, y axis) after 10 doublings. Significant genes (LFC > 0.75 and FDR < 0.1) are highlighted in green. Genes in the top right quadrant increase the expected fitness. RNF10 is color-coded in magenta. **(B)** Primer extension analysis of orthogonal 18S rRNA (WT or A1824C) upon siRNA knockdown of RNF10. Bottom: Immunoblots for RNF10 and β-actin. **(C)** Quantification of primer extension analysis shown in (B). 18S:WT levels with non-targeting siRNA was set to 1. Significance was calculated using Student’s t-test (***: p < 0.001, **:p < 0.01). **(D)** Primer extension analysis of orthogonal 18S rRNA (WT or A1824C) in parental HeLa and GCN2KO cells upon RNF10 knockdown by siRNA. Bottom: Immunoblots for GCN2, RNF10, and β-actin. **(E)** Quantification of primer extension analysis shown in (D) (n= 3). Mut: A1824C. RNF10 knockdown in GCN2KO background showed no further increase (yellow and orange bars). Student’s t-test, ns: p > 0.05. **(F)** Immunoblots for uS3 ubiquitination status from affinity-purified wild-type and decoding-incompetent (Mut, A1824C) ribosomes in parental HeLa and RNF10KO cells, analyzed by indicated antibodies. Ribosomal protein eS7 serves as a loading control in purified samples. Ub denotes ubiquitinated uS3. Molecular weights of the markers (kDa) are indicated. **(G)** Similar to (F), immunoblots for uS3 ubiquitination status in parental HeLa and GCN2KO cells. Reduction in uS3 ubiquitination in GCN2KO cells are quantified in Figure S6E.

Although recent studies have implicated RNF10 in recognizing stalled 43S PIC and 80S ribosomes^39,40^, its connection with GCN2 in 18S NRD had not been explored. We thus began by confirming the growth phenotype. As expected, RNF10 ablation in GCN2KO cells expressing 18S:A1824C rRNA did not further exacerbate cell fitness (Figure S6D). We next tested the effect of RNF10 on nonfunctional 18S rRNA turnover. Upon siRNA-mediated knockdown of RNF10 in the parental background, we observed a ∼4-fold increase in 18S:A1824C rRNA levels compared to the non-targeting knockdown (Figures 6B and 6C), phenocopying the observed effects upon GCN2 ablation (Figures 3A and 3B). Our genetic interaction screen revealed a buffering effect (i.e., positive genetic interaction) (Figure S6C) between GCN2 and RNF10 (Figure 6A), suggesting that GCN2 and RNF10 likely lie within the same pathway in degrading nonfunctional 18S rRNA. Therefore, we carried out an epistasis experiment by knocking down RNF10 in parental HeLa cells and GCN2KO cells that express 18S:WT or 18S:A1824C rRNA. If our model that GCN2 and RNF10 lie within in the same pathway is correct, then we should not observe an additive effect upon GCN2 and RNF10 double ablation. GCN2 knockout stabilized 18S:A1824C rRNA as shown before (Figures 6D, 6E, and S4A) and RNF10 ablation in GCN2KO cells did not further stabilize 18S:A1824C rRNA levels (Figures 6D and 6E), consistent with the genetic interaction screen. Taken together, these results demonstrate a genetic and functional epistatic relationship between GCN2 and RNF10 in promoting nonfunctional 18S rRNA turnover.

### Ubiquitination of decoding-incompetent ribosome is dependent on GCN2-RNF10 axis

Recent studies revealed RNF10 ubiquitinates ribosomal protein uS3 to trigger the degradation of 40S ribosomal proteins^39,40,63^. Indeed, purified decoding-incompetent ribosomes exhibit high levels of mono-ubiquitinated uS3 compared to wild-type ribosomes (Figure 6F, lanes 1 and 2). Notably, RNF10 knockout abolishes uS3 ubiquitination (Figure 6F, lanes 3 and 4), consistent with its reported role in targeting stalled 43S PIC and 80S ribosomes for degradation. Given the epistatic effect between GCN2 and RNF10 (Figures 6D and 6E), we next evaluated whether GCN2 impacts RNF10-dependent uS3 ubiquitination of decoding-incompetent ribosomes. Strikingly, GCN2 knockout strongly reduced uS3 ubiquitination (Figures 6G and S6E), phenocopying the observed effect in RNF10KO cells (Figure 6F). These results provide support for the function of GCN2 in triggering 18S NRD by regulating RNF10-mediated uS3 ubiquitination of decoding-incompetent ribosomes.

## DISCUSSION

Our work provides compelling evidence for the conservation of 18S NRD in mammalian cells and reveals a negative-feedback regulatory mechanism by which cells finetune translation initiation to surveil defective ribosomes engaged in translation, resulting in their elimination. Specifically, expression of the nonfunctional 18S:A1824C rRNA results in decoding-incompetent ribosomes arrested at start codons. Such ribosome stalls at translation start sites effectively trigger two distinct molecular events— GCN2-dependent eIF2⍺ phosphorylation and RNF10-dependent ubiquitination of the stalled ribosomes, both of which are required for elimination of nonfunctional 18S rRNA. GCN2 activation by arrested decoding-incompetent ribosomes triggers eIF2⍺ phosphorylation and a subsequent pro-survival dampening of global translation initiation, enabling RNF10 to detect and flag decoding-incompetent ribosomes for destruction (Figure 7A, top). The inverse correlation between the elevated levels of 43S-80S collisions at translation start sites (Figure 5F) and the reduction in RNF10-mediated ubiquitination of decoding-incompetent ribosomes (Figure 6G) in GCN2KO cells can be explained by a model where initiation collisions between scanning 43S PICs and decoding-incompetent ribosomes occludes RNF10, thereby impeding 40S subunit degradation and nonfunctional 18S rRNA turnover (Figure 7A, bottom).

**Figure 7.**
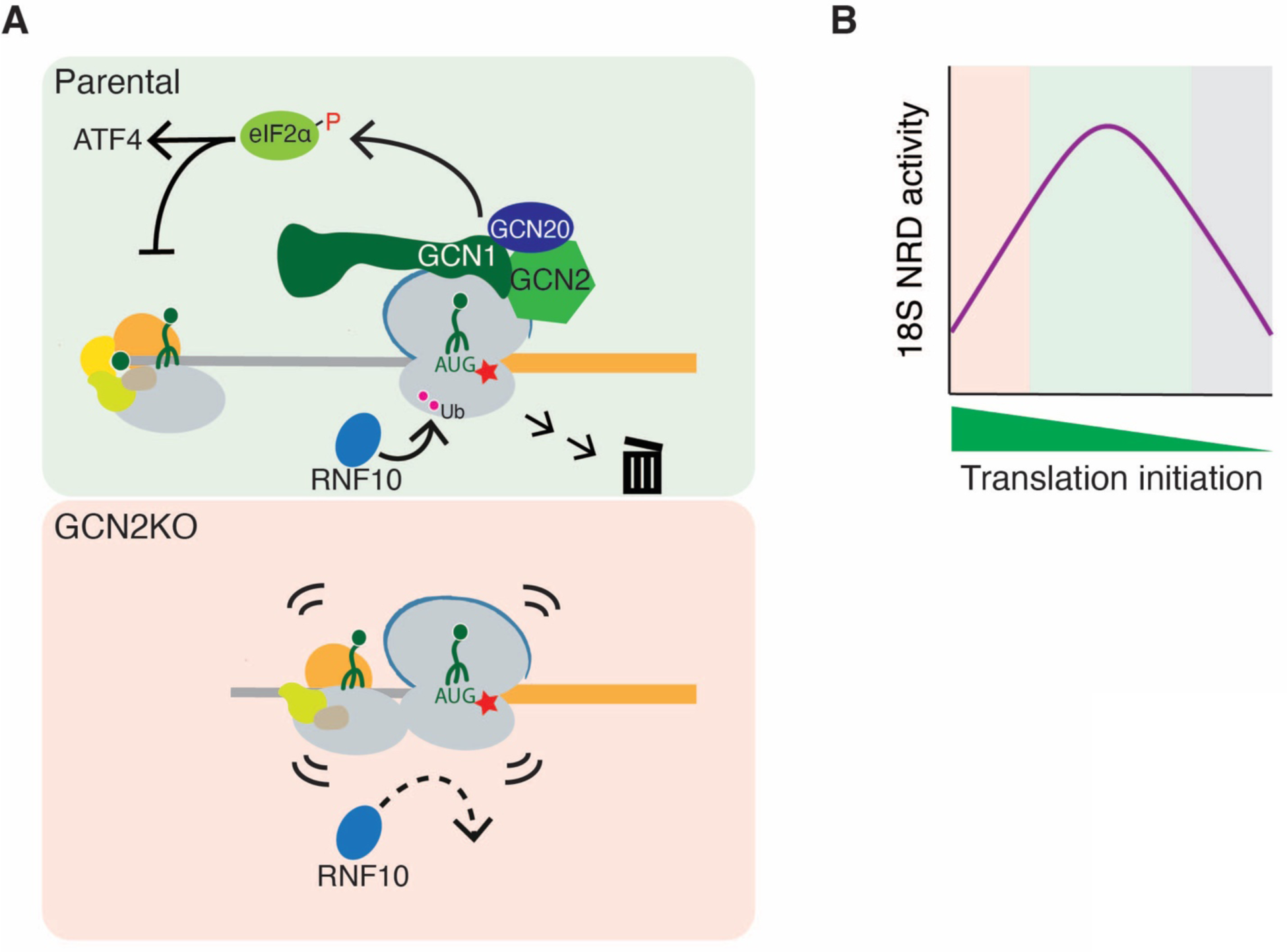
Working model for the ISR-regulated ribosome functionality surveillance. **(A)** Proposed working model for ribosome functionality surveillance elicited by RNF10 and the GCN complex. Red star denotes the decoding base mutation (A1824C). **(B)** Schematic for optimal translation initiation being critical for nonfunctional 18S rRNA turnover. 18S NRD activity requires optimal levels of translation initiation (green) and is dampened when initiation is unchecked (red) or completely inhibited (grey).

Our findings indicate that intermediate levels of phosphomimetic eIF2⍺^S51D^, as opposed to high levels, effectively complement the loss of GCN2 in degrading nonfunctional 18S rRNA (Figures 4E-4G), suggesting that 18S NRD activity requires optimal levels of translation initiation (Figure 7B). We propose that such levels of translation initiation are maintained by GCN2-mediated eIF2⍺ phosphorylation to prevent 43S-80S collisions. This negative feedback regulation enables RNF10 to effectively surveil stalled ribosomes (Figure 7B, green box). In the absence of the translation initiation brake mounted by GCN2, unchecked initiation results in 43S-80S collisions, allowing defective ribosomes engaged in translation to evade the quality surveillance (Figure 7B, red box). Lastly, high levels of phosphomimetic eIF2⍺^S51D^ deplete the eIF2 ternary complex and reduce 43S PIC formation and translation initiation (Figure 7B, grey box). Therefore, decoding-incompetent ribosomes are not engaged in a “test run” to examine their functionality. Our model is parsimonious and easily reconciles our findings with an earlier observation that ongoing translation is required for 18S NRD in budding yeast^3^. In this view, the GCN complex effectively functions as a sensor for stalled ribosomes with a vacant A site, enabling the negative feedback loop to regulate ribosome functionality surveillance.

In experiments aimed to address how decoding-incompetent ribosomes activate GCN2-dependent eIF2⍺ phosphorylation, we found that the decoding-inactivating mutation (A1824C) renders the ribosomal A site empty (Figure 4C), a crucial prerequisite for activation of GCN2 independently of uncharged-tRNA as reported recently^18,24^. Additionally, GCN1 is preferentially recruited to arrested decoding-incompetent ribosomes (Figure 4D). In contrast to the translational perturbations utilized in numerous previous studies, including translation elongation inhibitors, alkylating agents that damage mRNAs, and depletion of tRNA or termination factor, all of which could lead to ribosome collisions during elongation, decoding-incompetent ribosomes are predominantly arrested at start codons throughout the transcriptome (Figures 1F-1H) without inducing detectable 80S-80S ribosome collisions in the 5ʹ UTRs (Figure S5E). We show that such prolonged ribosome arrest activates the GCN2-dependent ISR. These results explain numerous recent findings. First, GCN2 is preferentially activated by ribosome collisions with an empty A site in the leading ribosomes^18,24^. Second, GCN1 associates not only with collided ribosomes but also with stalled monosomes at nonoptimal codons in the 3ʹ UTRs upon induced readthrougth^17,18,56,64^. Third, GCN1/20 co-sediments with monosomes regardless of amino acid starvation or stress conditions that induce ribosome collisions^59,65^. Integrating our results with recent studies, we speculate that GCN1 is recruited to stalled monosomes, thereby signaling GCN2 to activate the ISR before trailing ribosomes collide. Our work extends these studies by showing that GCN1 facilitates the activation of GCN2 potentially through its association with arrested decoding-incompetent ribosomes independently of ribosome collisions, a mechanism that likely works in concert with uncharged-tRNA dependent GCN2 signaling.

Collided ribosomes trigger both the GCN2-dependent ISR and the ZAK⍺-dependent RSR^17^ whereas our data indicate stalled elongating monosomes can only activate the GCN2-mediated ISR. Given the temporal order of ribosome collisions on mRNAs— where ribosomes stall first, and trailing ribosomes catch up and collide— it makes considerable sense that the pro-survival ISR can be initiated as early as translational perturbation (i.e., ribosome stalling) occurs. This activation of the ISR is sustained through subsequent ribosome collisions, effectively dampening translation initiation and promoting cell survival. These results highlight a mechanism where a measured and well-timed stress response pathway maintains translational homeostasis. The fact that GCN2 is typically found in sub-stoichiometric levels compared to GCN1/20 across various cell types^66^ implies a multifaceted role for GCN1/20 beyond its function in the GCN2-dependent ISR. Indeed, recent studies have uncovered that GCN1 is involved in sensing translational perturbations to facilitate ribosome-mediated mRNA decay and proteolytic degradation of translation factors^67–70^. Future structural and biophysical studies of the relationship between the GCN2/1/20 complex and the ribosome (stalled vs. collided) will provide spatially and temporally resolved insights into how these intricate regulatory mechanisms are coordinated.

While there is evidence indicating cap-tethered 43S PIC scanning limits ribosome loading and hence 43S-80S collisions at start codons^71^, it is possible that mRNA caps become available to eIF4E given the long half-life of nonfunctional 18S rRNA (Figure 1E) and potentially prolonged ribosome arrest at start codons in the cell (Figure 1F). We implemented the recent improvement in ribosome profiling^52^ by using nuclease P1 to capture 43S-80S footprints at translation start sites induced by decoding-incompetent ribosomes (Figures 5C-5F). These findings are further supported by several lines of evidence confirming that the isolated footprints are derived from 43S-80S rather than disomes (80S-80S). First, nonfunctional 18S rRNA does not activate ZAK⍺ and the downstream RSR (Figure S5D), a hallmark of ribosome collisions during translation elongation^17^. Second, disome profiling does not show detectable 80S-80S collisions at start codons even in GCN2-ablated cells expressing nonfunctional 18S rRNA (Figure S5E). Footprints protected by 43S-80S collision complexes (∼57 nt in length) are readily discernible from typical disome footprints (∼72 nt in length)^52^, and this distinction will enable future efforts to explore the potential role of initiation collisions in the regulation of protein synthesis.

Notably, our model where 43S-80S initiation collisions impede the ability of RNF10 to ubiquitinate uS3 (Figure 7A) aligns with a recent finding that Fap1, the yeast ubiquitin E3 ligase responsible for elongating the ubiquitin chain on uS3, selectively targets stalled individual ribosomes but not collided ribosomes (i.e., disomes) for elimination^6^. While the ribosomal protein uS3 does not appear to be polyubiquitinated in mammalian cells (^38–40^ and Figures 6F and 6G), it is possible that RNF10 regulates the substrate specificity of the 18S NRD quality control pathway for stalled monosomes rather than collided ribosomes. Overall, our finding that RNF10 flags decoding-incompetent ribosomes for elimination is broadly consistent with prior studies of RNF10 in mammals^39,40^ and Mag2 in yeast^5,6^. Given that both initiation and elongation inhibitors trigger uS3 ubiquitination^39,40^, further investigation will reveal how RNF10 surveils its biological targets (i.e., nonproductive 43S complex and stalled 80S) and directs them toward degradation.

Importantly, ribosome collisions activate RQC to monitor mRNAs and nascent polypeptide chains whereas ribosome stalling appears to trigger a surveillance pathway to interrogate the functionality of stalled individual ribosomes. As molecular mechanisms associated with these surveillance pathways are just beginning to be unraveled, our results and approach provide a framework for understanding the regulation of stalled elongating ribosomes prior to the occurrence of ribosome collisions.

## ACKNOWLEDGMENTS

We would like to thank Niladri Sinha and Tom Dever for critical reading of the manuscript; Johnathan Bohlen for advice on translation complex profile sequencing; Jeff Carrell and Megan Karwan at the NCI-Frederick Flow Cytometry Core Facility for assistance in cell sorting. This work was supported by the Intramural Research Program of the National Institutes of Health, National Cancer Institute, Center for Cancer Research (1ZIABC012037).

## AUTHOR CONTRIBUTIONS

Conceptualization, A.R.C. and C.C.C.W.; Methodology, M.S., A.S., I.M.S. J.T.M., and C.C.C.W.; Investigation, A.R.C., A.S., M.S., S.J.T., I.M.S., and C.C.C.W.; Resources, J.T.M.; Visualization, A.R.C., C.C.C.W., and W.G.; Data Curation, W.G.; Writing, A.R.C. and C.C.C.W.; Supervision, C.C.C.W.

---

**Figure S1.**
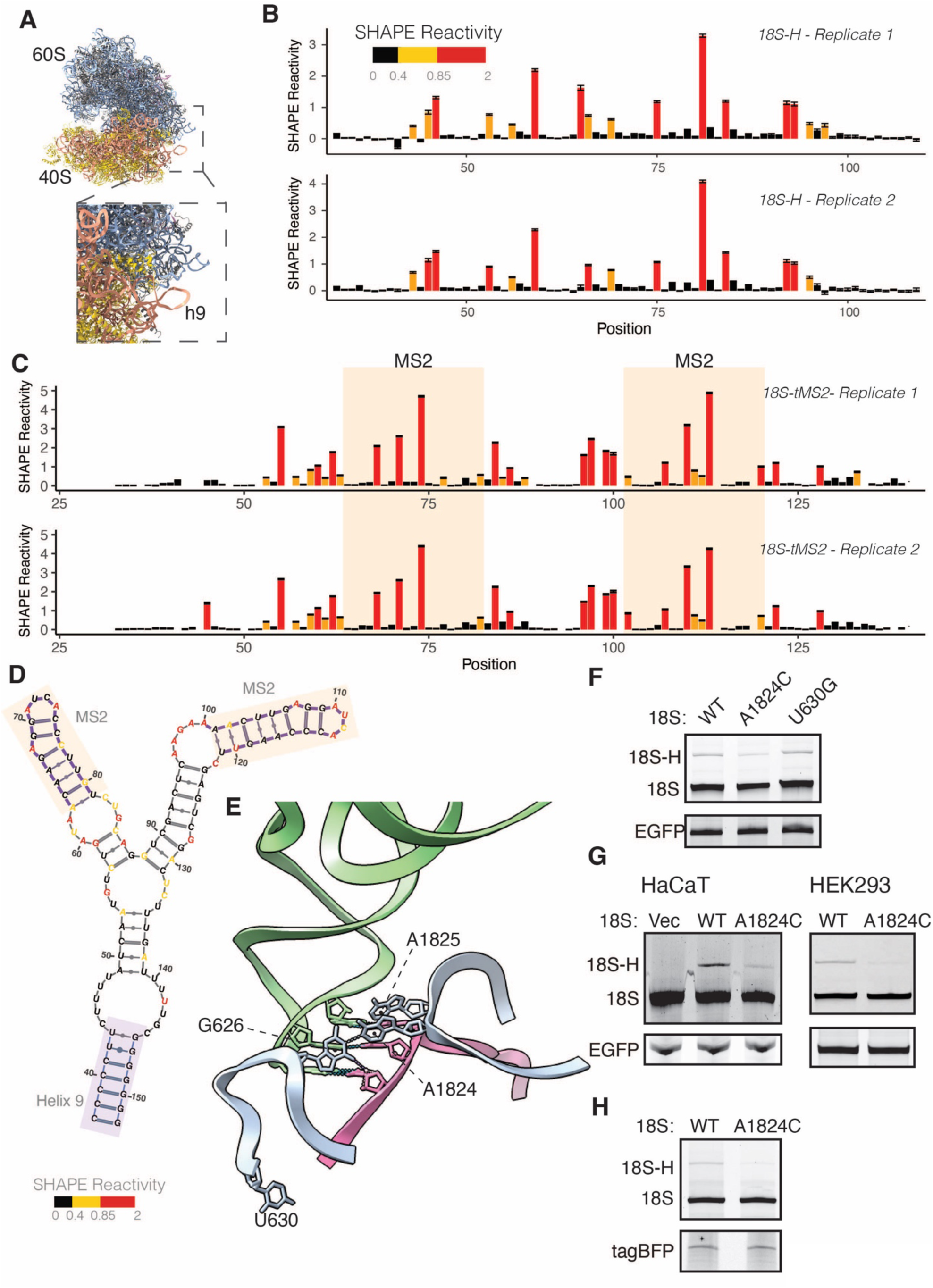
Establishing an orthogonal expression system to study nonfunctional 18S rRNA turnover, related to Figure 1. **(A)** Cryo-EM structure of human ribosome with the ribosomal 40S and 60S subunits denoted. Inset: a close-up view of 18S rRNA helix 9 where an exogenous tag was inserted. **(B)** *In vivo* SHAPE derived reactivities from orthogonal 18S expression system harboring a hybridization (18S-H) tag in helix 9 probed with 5NIA. Bars represent SHAPE reactivities at single nucleotide resolution. Nucleotides exhibiting high (2- 0.85), medium (0.85- 0.4), and low (0- 0.4) SHAPE reactivity are color-coded in red, yellow, and black, respectively. Two biological replicates are shown. **(C)** Similar to (B), *in vivo* SHAPE derived reactivities from the orthogonal 18S expression system harboring tandem MS2 stem loops (tMS2) in helix 9 probed with 5NIA. MS2 stem loops are shaded in orange **(D)** SHAPE informed minimum free energy secondary structure of tMS2 insertion at the tip of 18S helix 9. Each nucleotide is color-coded by SHAPE reactivity. **(E)** Cryo-EM structure of human ribosome depicting the decoding center in the 40S subunit (PDB: 8g6j). The monitoring bases (G626, A1824, and A1825) and U630 that has no role in decoding are denoted. tRNA and mRNA are shown in green and pink, respectively. **(F)** Primer extension analysis of 18S rRNA carrying indicated mutation. WT: wild-type. A1824C: decoding center mutation. U630G: a benign mutation. EGFP was used as a transfection and loading control. Also see Figure 1C for corresponding Northern blotting. **(G)** Primer extension analysis of 18S rRNA carrying A1824C mutation in human keratinocyte cells (HaCaT) and HEK293 cells, validating the conservation of 18S NRD among different human cells. **(H)** Primer extension analysis of 18S rRNA levels form HeLa cells that constitutively express 18S:WT or A1824C. tagBFP serves as a control for integration and loading.

**Figure S2.**
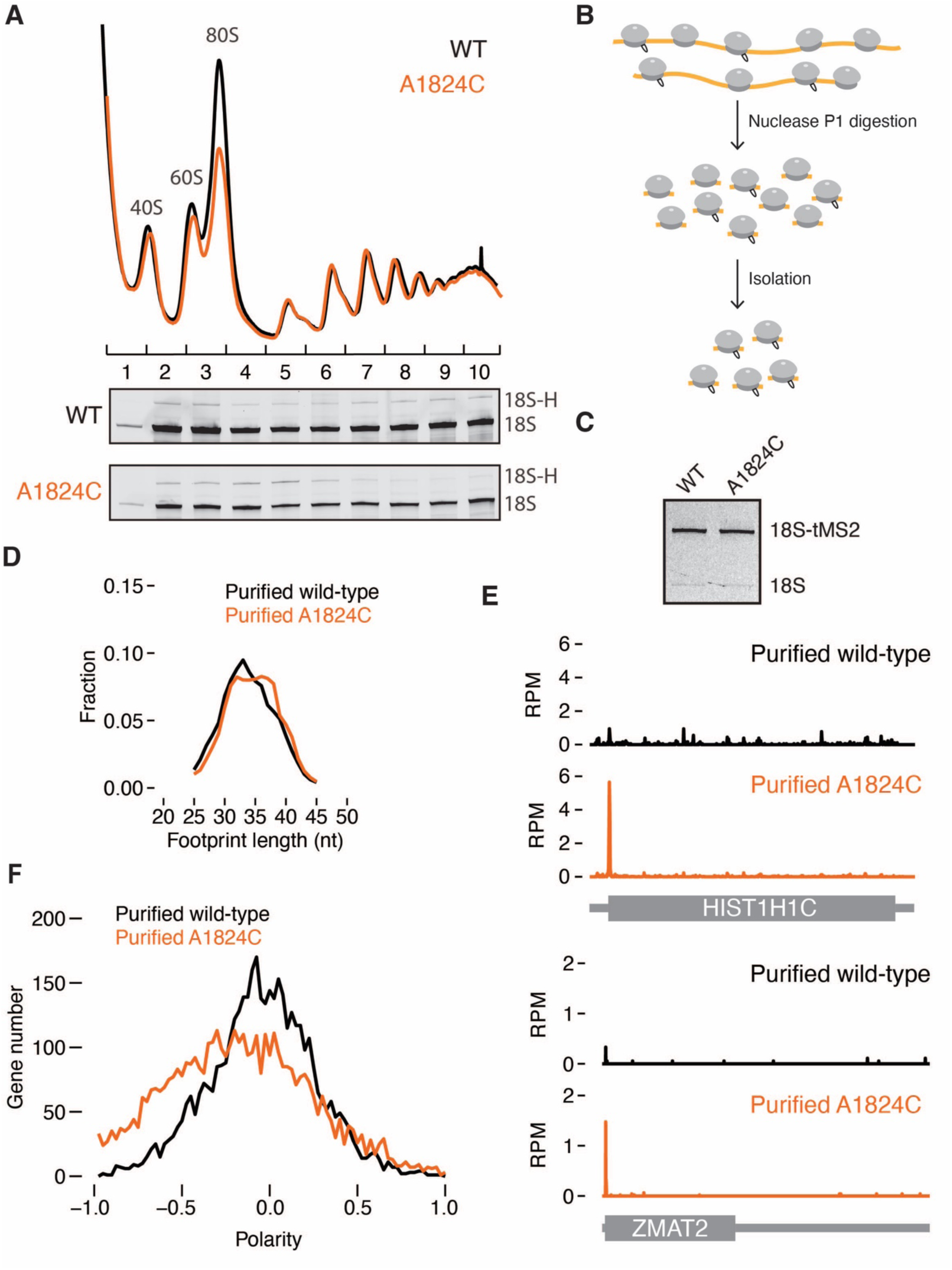
18S:A1824C rRNA is incorporated into the translating ribosome pool, related to Figure 1. **(A)** Representative polysome profiles over a 10-50% sucrose gradient from whole cell lysates of cells expressing 18S:WT (black) or 18S:A1824C (orange). Bottom: Primer extension analysis of rRNA extracted from respective fractions. **(B)** Schematic for affinity purification of tMS2-tagged ribosomes. To avoid cleaving inserted tMS2 tag in 18S rRNA, lysates were partially digested with nuclease P1 and incubated with MS2 coat protein to isolate tagged ribosomes. **(C)** Primer extension analysis validated the enrichment of tMS2-tagged ribosomes (top band) from endogenous ribosomes (bottom band). **(D)** Size distribution of monosome footprints affinity-purified wild-type (black) and decoding-incompetent ribosomes (A1824C, orange). **(E)** Enrichment of decoding-incompetent ribosome footprints at start codons of two representative transcripts, HIST1H1C and ZMAT2. **(F)** Polarity score calculating ribosome occupancy across ORFs from affinity-purified wild-type (black trace) and decoding-incompetent ribosomes (A1824C, orange trace).

**Figure S3.**
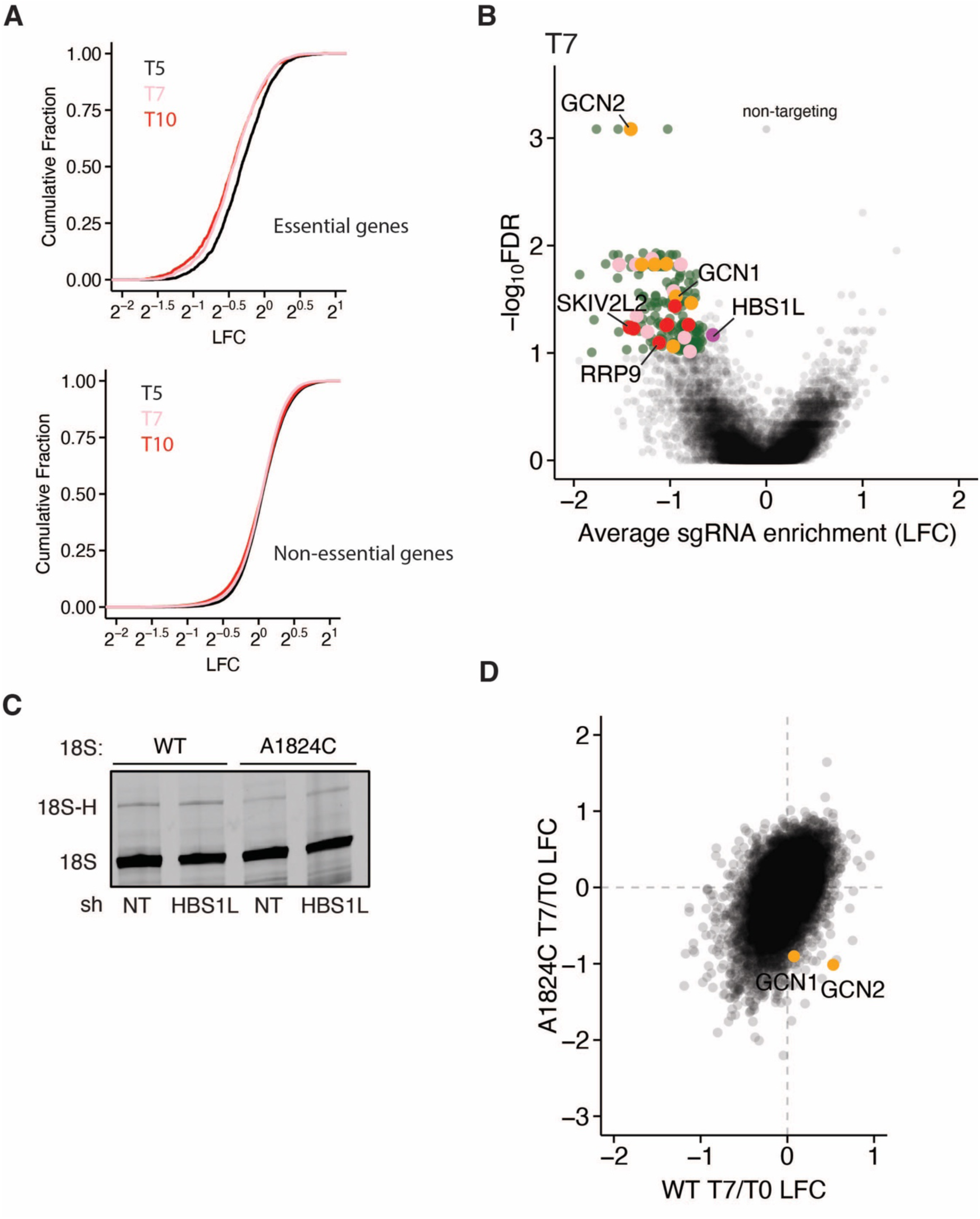
A Fitness-based CRISPR screen to identify factors involved in 18S NRD, related to Figure 2. **(A)** Cumulative distribution of changes in sgRNAs targeting essential genes (top) or non-essential genes (bottom) at T5, T7, and T10 relative to T0. LFC: log_2_ fold change. **(B)** Volcano plot showing average sgRNA enrichment (log_2_ fold change [LFC], x axis) versus false-discovery rate (FDR) (- logFDR, y axis) at T7. Significant genes (LFC < 0.75 and FDR < 0.1) are highlighted in green. Color of dots corresponds to the enriched Gene Ontology (GO) analysis shown in Figure 2D. HBS1L is shown in purple. Also see Figure 2C. **(C)** Primer extension analysis of orthogonal 18S rRNA (WT or A1824C) in cells depleted of HBS1L by shRNA. **(D)** Scatter plot showing de-enrichment of GCN2 and GCN1 sgRNAs after 7 doubling by taking the ratio of T7 over T0 in cells expressing 18S:WT (X axis) or 18S:A1824C cells (y-axis).

**Figure S4.**
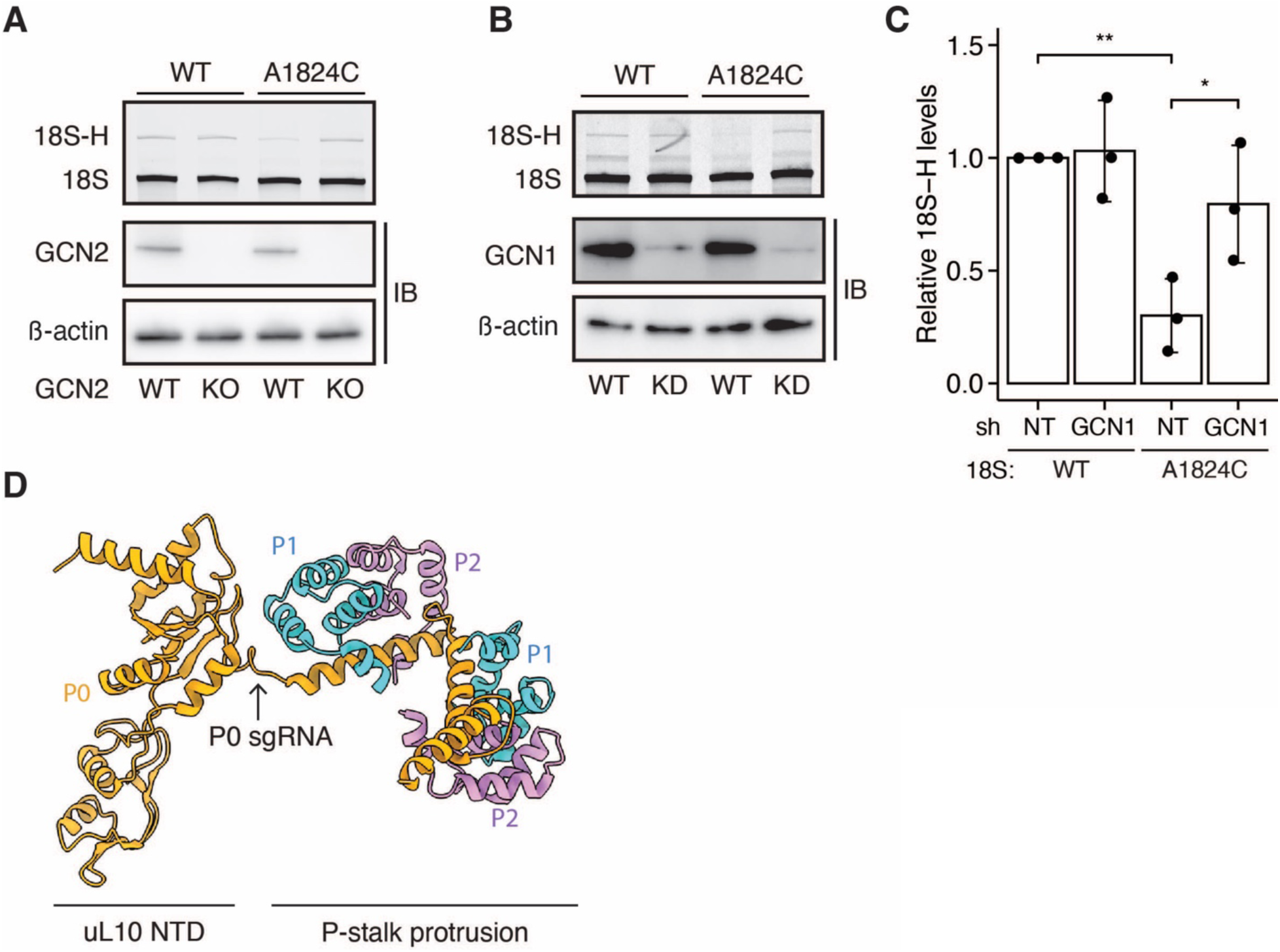
GCN complex and ribosomal P-stalk promote 18S NRD, related to Figure 3. **(A)** Primer extension analysis of orthogonal 18S rRNA (WT or A1824C) in GCN2KO cells. Bottom: Control immunoblots for GCN2 and β-actin. **(B)** Representative primer extension analysis of orthogonal 18S rRNA upon shRNA-mediated knockdown of GCN1. Bottom: Immunoblots for GCN1 and β-actin. **(C)** Quantification of primer extension analysis upon GCN1 knockdown in (B) (n= 3). Significance was calculated using unpaired Student’s t-test (**: p < 0.01, *: p < 0.05). **(D)** Cryo-EM structure of the ribosomal P-stalk (PDB: 4v6x). The uL10 (P0) sgRNA used in this study targets the helical region joining uL10 NTD with the P-stalk protrusion. Ribosomal proteins P0, P1, and P2 are color-coded to show the heteropentameric structure.

**Figure S5.**
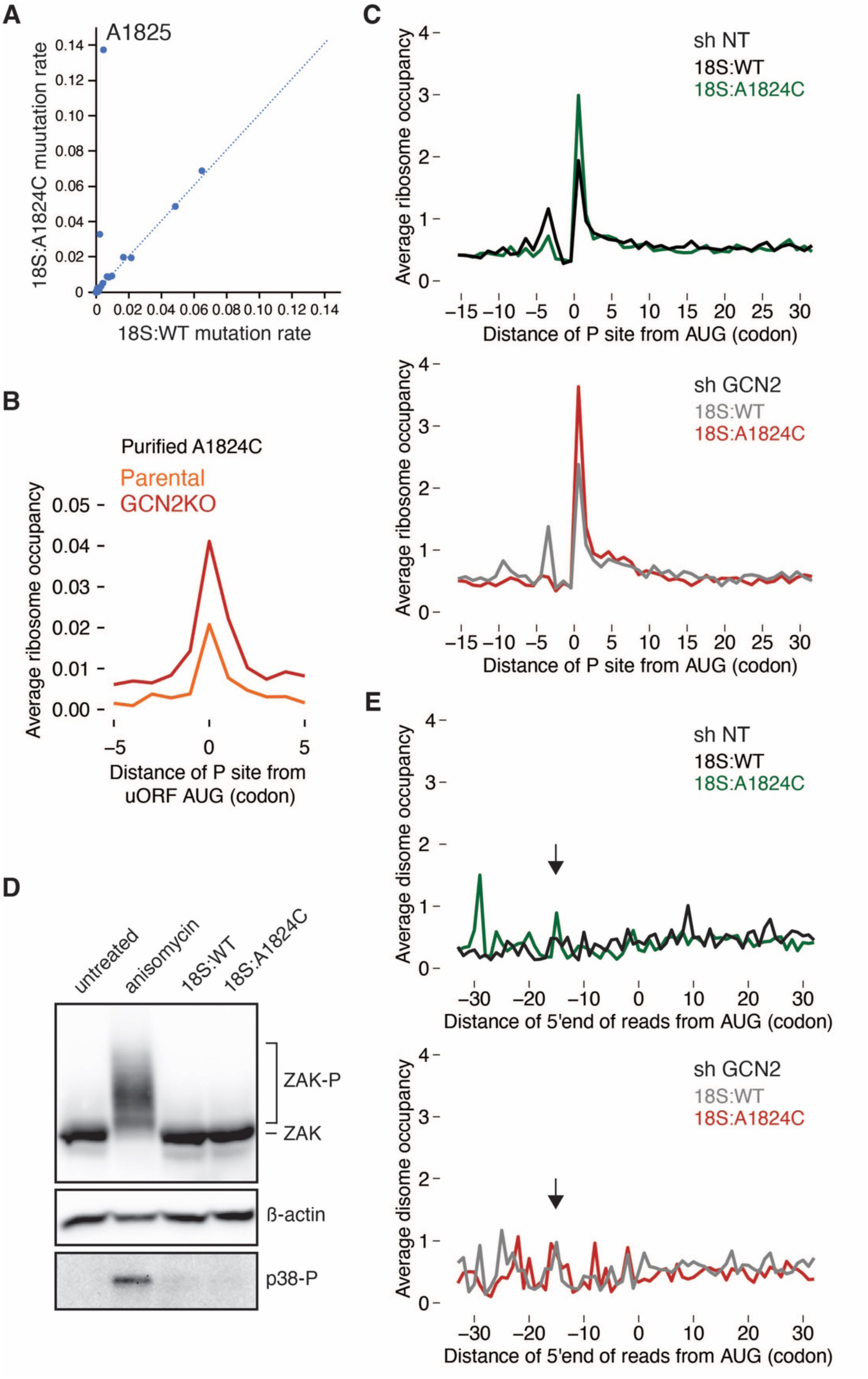
18S:A1824C rRNA renders the ribosomal A site vacant but does not result in ribosome collisions, related to Figures 4 and 5. **(A)** Scatter plot comparing DMS mutation rates of nucleotides located within the ribosomal A site of wild-type and decoding-incompetent (A1824C) ribosomes, biological replicate 2. Also see Figure 4C. **(B)** Average ribosome occupancy of affinity-purified decoding-incompetent ribosomes aligned at annotated uORF start codons from parental HeLa (orange trace) and GCN2KO (red trace) cells. Offsets were set to show the ribosomal P site of the footprints. **(C)** Metagene analysis of average ribosome occupancy aligned at start codons. Offsets were set to the P site of the ribosome footprints. Top: Ribosome footprints derived from 18S:WT (black trace) and 18S:A1824C cells (green trace) expressing non-targeting shRNA. Bottom: Ribosome footprints derived from 18S:WT (grey trace) and 18S:A1824C cells (red trace) expressing GCN2 shRNA. **(D)** Immunoblots for ZAK⍺ and p38 phosphorylation in GCN2KO cells treated with an intermediate dose of anisomycin that triggers ribosome collisions (at 0.1 µg/mL for 30 min) or transfected with a plasmid encoding 18S:WT or 18S:A1824C rRNA compared to untreated cells. ZAK⍺ phosphorylation is monitored through Phos-tag gels that retard the migration of phosphorylated proteins. p38 phosphorylation is monitor through a phospho-p38 antibody. β-actin is a loading control. Neither 18S:WT or 18S:A1824C induces ZAK⍺ phosphorylation, indicating the absence of ribosome collisions. **(E)** Average disome occupancy aligned at start codons. 5ʹ ends of disome footprints are shown. Top: Disome footprints derived from 18S:WT (black trace) and 18S:A1824C cells (green trace) expressing non-targeting shRNA. Bottom: Disome footprints derived from 18S:WT (grey trace) and 18S:A1824C cells (red trace) expressing GCN2 shRNA. Arrows indicate the anticipated positions if ribosome collisions (80S-80S) were to occur.

**Figure S6.**
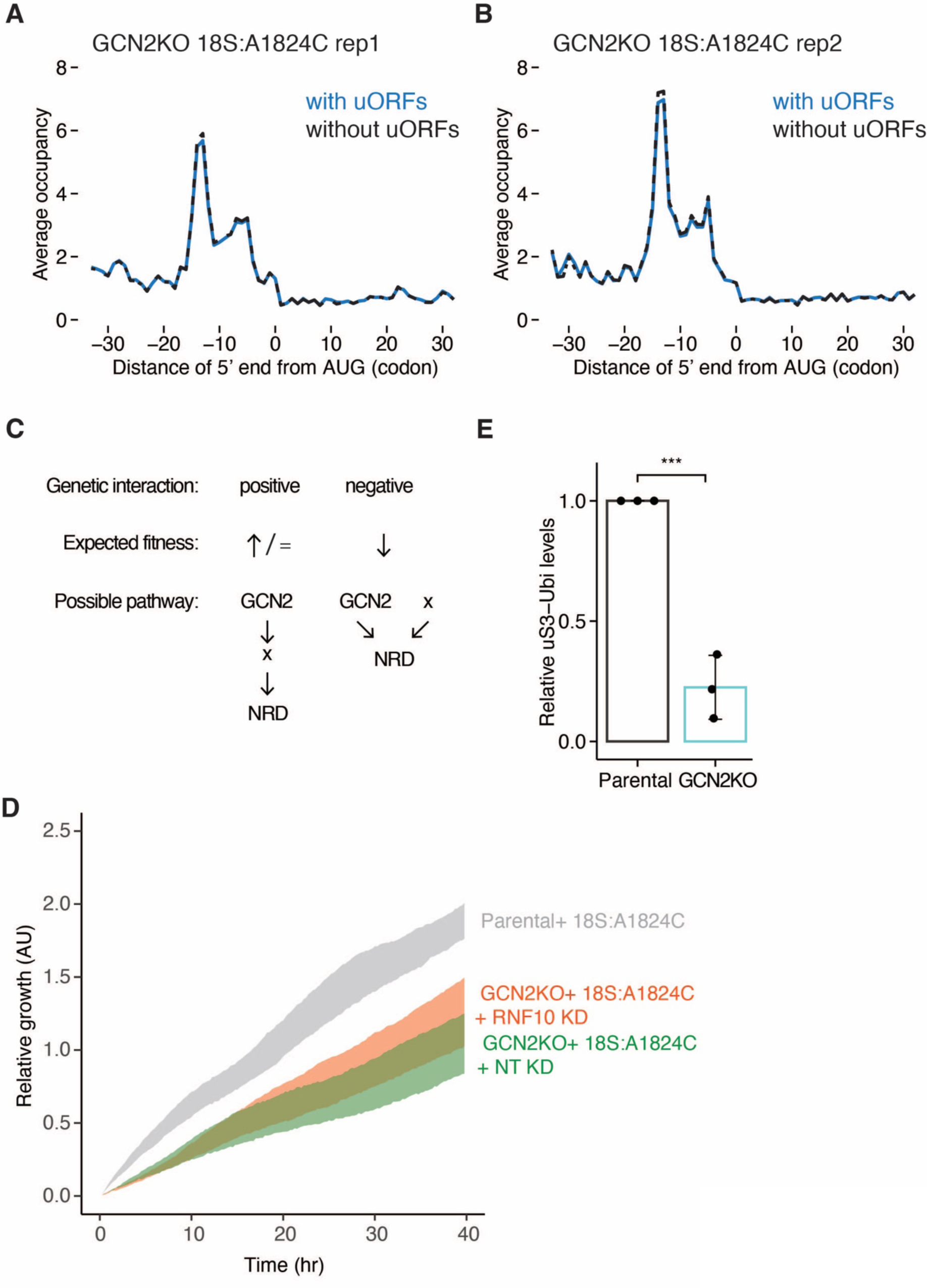
Loss of GCN2 induces 43S-80S collisions at translation start sites and blocks RNF10-mediated ubiquitination of ribosomal protein, related to Figures 5 and 6. **(A-B)** Metagene analysis of average 43S-80S occupancy aligned at start codons. 5’ ends of the footprints are shown relative to the start codon. The 43S-80S footprints are derived from affinity-purified decoding-incompetent ribosomes (18S:A1824C) in GCN2KO cells, as shown in Figure 5F. This analysis was performed including transcripts with uORFs (blue trace) or without uORFs (black trace). **(C)** Expected fitness and possible biological pathways resulting from ablation of x gene in combination of GCN2KO. **(D)** Quantification of cell growth by xCELLigence (x axis). Growth phenotype was compared across three cell lines: parental HeLa cells expressing 18S:A1824C rRNA (grey) and GCN2KO cells expressing 18S:A1824C rRNA with nontargeting (green) or RNF10 shRNA (orange). **(E)** Quantification of uS3 ubiquitination by immunoblotting (n= 3) as shown in Figure 6G from purified decoding-incompetent ribosomes (18S:A1824C) in parental HeLa and GCN2KO cells. Significance was calculated using Student’s t-test (***: p < 0.001).

